# Proterozoic Tectonic Drivers Underpin Flavobacterial Diversification

**DOI:** 10.1101/2025.04.01.646730

**Authors:** Hao Zhang, Haiwei Luo

**Affiliations:** Key Laboratory of Quantitative Synthetic Biology, Shenzhen Institute of Synthetic Biology, Shenzhen Institute of Advanced Technology, Chinese Academy of Sciences, Shenzhen, China; Simon F. S. Li Marine Science Laboratory, School of Life Sciences and State Key Laboratory of Agrobiotechnology, The Chinese University of Hong Kong, Shatin, Hong Kong SAR; Department of Earth and Environmental Sciences, The Chinese University of Hong Kong, Shatin, Hong Kong SAR; Institute of Environment, Energy and Sustainability, The Chinese University of Hong Kong, Shatin, Hong Kong SAR

**Author notes:** Corresponding author: Haiwei Luo.

**Keywords:** Flavobacteria, molecular dating, Columbia supercontinent, niche transition, diversification rate

## Abstract

Flavobacteria are keystone taxa in global carbon cycling, degrading complex glycans in marine and terrestrial ecosystems. In both environments, polysaccharides constitute a major fraction of organic matter but differ in origin: marine glycans primarily derive from micro- and macroalgae, while terrestrial counterparts originate from land plants. Flavobacteria deploy distinct suites of carbohydrate-active enzymes (CAZymes) tailored to these habitat-specific substrates, yet the evolutionary drivers of their diversification remain unresolved. Two competing hypotheses exist: one posits glycan specialization as the primary driver of divergence, while the other implicates extrinsic geological factors in shaping their evolution. By integrating mitochondrial- and plastid-based molecular clocks with eukaryotic fossil calibrations, we infer that flavobacteria emerged between 2.15 and 1.98 billion years ago (Gya), shortly after the Great Oxidation Event (GOE; 2.4-2.32 Gya). Subsequent diversification involved three independent marine-to-non-marine transitions during the Proterozoic Eon (1.98-1.70 Gya, 1.72-1.40 Gya, and 1.28-1.14 Gya), temporally aligned with the formation and the fragmentation of the Columbia supercontinent and preceded the evolution of the major glycan contributors in both marine and non-marine niches. This temporal mismatch disfavors the glycan specialization hypothesis, instead implicating tectonic-driven habitat shifts as the primary driver of lineage diversification. Non-marine flavobacteria exhibited higher turnover but lower net diversification rates than marine counterparts, reflecting the challenges of adapting to fragmented non-marine niches. Genome erosion and deleterious mutation accumulation further constrained reverse transitions, locking lineages into non-marine habitats. Our findings highlight Proterozoic tectonics, rather than substrate-specific CAZyme innovation, as the catalyst for flavobacteria’s evolutionary success across Earth’s carbon-rich biomes.

## Introduction

Flavobacteria, a major deep-branching lineage within the phylum Bacteroidetes (here referring to the assemblage of *Flavobacteriaceae* and *Weeksellaceae* per Zhang et al. 2019), are ubiquitous marine and non-marine microbes renowned for their exceptional ability to degrade complex polysaccharides (glycans). This capability positions them as keystone taxa in global carbon cycling (Kirchman, 2002). In marine ecosystems, flavobacteria dominate during phytoplankton blooms, hydrolyzing sulfated algal glycans such as ulvan and laminarin through specific glycoside hydrolases and sulfatases (Teeling et al., 2012; Teeling et al., 2016). These enzymatic processes release dissolved organic carbon (DOC) into the microbial loop, fuelling 15-30% of marine heterotrophic activity (Moran et al., 2022). Likewise, in terrestrial and freshwater habitats, flavobacteria decompose plant-derived lignocellulose and pectin through glycoside hydrolases and polysaccharide lyases, facilitating soil organic matter turnover and rhizosphere nutrient cycling (Kolton et al., 2013).

Genomic analyses have revealed repeated marine-to-non-marine transitions in flavobacterial evolution, marked by dynamic gene gain/loss events linked to habitat adaptation (Zhang et al., 2019). However, whether these transitions were driven by substrate competition (e.g., glycan resource partitioning) or extrinsic geological factors remains unresolved. A central challenge lies in establishing a robust timeline for flavobacterial evolution due to the great scarcity of bacterial fossil records for molecular clock analysis. Unlike eukaryotes, whose fossil record extends back to the mid-Proterozoic eon (e.g., the Gaoyuzhuang Formation at 1.56 Gya and the Greyson Formation at 1.45 Gya) (Zhu et al., 2016; Adam et al., 2017), bacteria rarely preserve morphological signatures. While Cyanobacteria fossils have been used to constrain minimum ages for specific lineages, they provide limited temporal resolution for broader bacterial divergence events. Importantly, the lack of maximum age constraints for bacterial lineages can significantly impact the accuracy of molecular clock analyses, as maximum ages are critical for refining posterior age estimates (Hedges et al., 2018; Morris et al., 2018; Wang and Luo, 2021).

To address this, recent studies have used the extensive eukaryotic fossil record alongside mitochondria- and plastid-encoded genes derived from ancient endosymbiosis events involving Alphaproteobacteria and Cyanobacteria, respectively (Toft and Andersson, 2010; Perreau and Moran, 2022), to establish the robust temporal constraints for dating bacterial lineages (Sánchez-Baracaldo et al., 2017; Fournier et al., 2021; Wang and Luo, 2021; Zhang et al., 2023; Liao et al., 2024; Wang and Luo, 2025). These genes provide a molecular link between bacterial and eukaryotic lineages, allowing the transfer of well-constrained eukaryotic fossil calibrations to bacterial divergence events. By integrating these calibrations, the sequential dating approach allows for the accurate propagation of age information from eukaryotes to bacterial nodes, significantly improving the precision of divergence time estimates for bacterial lineages (Wang and Luo, 2021; Zhang et al., 2023; Liao et al., 2024). This approach bridges the bacterial-eukaryotic fossil divide, enabling the reconstruction of a robust evolutionary timeline for flavobacteria.

## Materials and Methods

### Taxonomy sampling and phylogenomic tree reconstruction

The genomic sequences of flavobacteria (including both the families *Flavobacteriaceae* and *Weeksellaceae*) used in this study were downloaded from the NCBI GenBank database (as of January 2023). We assessed the quality of the genomic assemblies in terms of completeness, contamination, and heterogeneity using the software CheckM v1.1.3 (Parks et al., 2015) with the taxonomy workflow and a manually assigned marker set specific to the class “Flavobacteriia.” The quality information derived from CheckM was then processed using the software dRep v3.4.2 (Olm et al., 2017) with the parameter “- -genomeInfo”. The average nucleotide identity (ANI) threshold was set to 0.7 (-pa 0.7), and the completeness and contamination thresholds were set to 90% and 10%, respectively (-comp 90 -con 10). For each genus within the phyla Bacteroidetes (excluding those belonging to the families *Flavobacteriaceae* and *Weeksellaceae*), Chlorobiia, and Cyanobacteria, a single representative species was randomly selected, with priority given to those marked with “reference” and “representative” and those in “complete” or “scaffold” level (Dataset S1). The open reading frames for these genomes were identified using the software Prodigal v2.6.3 (Hyatt et al., 2010). The predicted protein-coding genes were then annotated with the COG database (2020 updates) (Galperin et al., 2021) using the HMMER software package v3.3.2, with an e-value threshold of 1E-10 (Eddy, 2011). We extracted the sequences of 50 ribosomal genes across the bacterial species and aligned the sequences for each gene using the software MAFFT v7.520 (Katoh et al., 2002). The alignments were then trimmed using the software trimAl v1.4 (Capella-Gutiérrez et al., 2009) for concatenation. The phylogenomic tree of these bacterial species were reconstructed with IQ-Tree v2.2.6 (Minh et al., 2020). The best-fit amino-acid substitution model and rate heterogeneity model for each ribosomal gene partition were automatically determined by calling the ModelFinder program.

### Compiling molecular dataset for dating analysis

The marker genes for dating analysis were selected based on two considerations. First, genes that exhibit phylogenetic incongruence with the species tree due to horizontal gene transfer (HGT) or other evolutionary processes are unsuitable for molecular dating (Moody et al., 2022). Here, the verticality of marker genes were assessed using ΔLL as a proxy, which represents the difference in likelihood values between gene trees reconstructed with and without the backbone species tree (Moody et al., 2022). Second, genes exhibiting large evolutionary rate variations are more likely to violate the molecular clock and are difficult to correct with a relaxed clock model. Therefore, we assessed evolutionary rate differences between prokaryotes and eukaryotes using the Wilcoxon signed-rank test based on their branch lengths.

At the beginning of the study, we identified marker genes from three pre-compiled datasets: the 24 conserved mitochondrial genes shared between alphaproteobacterial and mitochondrial genomes (Wang and Wu, 2015), the 19 mitochondrion-originated genes likely transferred to eukaryotic nuclear genomes and conserved in bacterial genomes (Munoz-Gomez et al., 2019; Munoz-Gomez et al., 2022), and the 97 plastid marker genes showing no evidence of HGT among cyanobacterial species (Ponce-Toledo et al., 2017). For each dataset, we searched for marker genes in both eukaryotes and prokaryotes using BLASTP v2.6.0 with an e-value threshold of 1e-30. We then constructed the phylogeny for each marker gene using IQ-Tree v2.2.6 (Minh et al., 2020) with an auto-determined amino acid substitution model and a rate-heterogeneous model (Wang and Luo, 2021; Zhang et al., 2023).

Considering both factors, we ranked the ΔLL values (in increasing order) and the log-transformed p-value of the Wilcoxon signed-rank test (in increasing order) for each gene family, retaining those that ranked in the top 50% in both categories. As a result, we retained 13 plastid-encoded marker genes (Fig. S1), 9 mitochondrial-encoded marker genes, and 5 nuclear-encoded marker genes. Next, we manually examined the distribution patterns of bacterial and eukaryotic species in gene tree and removed those potentially derived from horizontal gene transfer. Following this curation, the nuclear-encoded and mitochondrial-encoded gene sets were excluded from further analysis, as using a small molecular dataset (fewer than five genes) can lead to biased posterior age estimates (Wang and Luo, 2024). The phylogeny of the 8 plastid-encoded marker genes that passed filtration is shown in Fig. S2.

### Implementation of Bayesian sequential dating strategy

The implementation of the Bayesian sequential approach divided the dating analysis into two steps. The first step, detailed in our recent study (Zhang et al., 2023), involved reconstructing the eukaryotic species tree using concatenated amino acid sequences of 50 plastid-encoded genes under the LG+G+C60 model. Posterior ages were then estimated using six non-redundant eukaryotic fossil-based calibrations (Table S1). The posterior ages of eukaryotic ancestors were then approximated to one of three distributions, including Gamma distribution, skew-normal distribution, and skew-t distribution, which were applied as the time prior in the second step dating analysis (Table S1; Fig. S3).

At the beginning of the second step of dating analysis, we grafted the species tree topology of eukaryotes (Archeaplastida) onto that of Cyanobacteria. The latter also includes Bacteroidetes and Chlorobiia. Using this synthetic species tree topology, we selected the best-fit clock model for each plastid-encoded marker gene via the stepping-stone method, implemented in the ‘mcmc3r’ program (dos Reis et al., 2018). Since calculating the marginal likelihood for concatenated sequences requires significant computational resources and time, we followed a recent approach (Wang and Luo, 2024) by calculating the likelihood separately for each marker gene. To meet the program’s requirements, we recoded the amino acid sequences into four nucleotide characters based on the Dayhoff4 recoding scheme. We found that seven of the plastid-encoded marker genes supported the auto-correlated rate model, while only one supported the independent rate model, indicating that the former is more suitable for this case (Table S2). Next, we calibrated the synthetic species tree topology using the approximated distributions derived from the first-step dating analysis (Zhang et al., 2023), as well as additional eukaryotic fossil-based calibrations, cyanobacterial fossil-based calibrations and the biomarker-based calibration (Table S1). The justification for these calibrations has been addressed in recent studies (Wang and Luo, 2021; Zhang et al., 2023; Wang and Luo, 2024).

The time-calibrated species tree and the marker genes were input into MCMCTree v4.10 (Yang, 2007) for posterior age estimation under the C60 model (Wang and Luo, 2024) (Fig. S4). To ensure MCMC convergence to the posterior distribution, we set the parameters “nsample” and “samplefreq” to 20,000 and 30, respectively, collecting a total of 600,000 samples for each run of the dating analysis. For each dating analysis, we plotted the correlation of the posterior mean ages between replicated runs. Convergence of the independent runs is achieved if the posterior ages closely align with the y=x line (Fig. S5).

### Assessing the reduction in effective population size in ancient time

By classifying non-synonymous substitutions into radical and conservative categories, we calculated the ratio of substitution rates between the two groups (i.e., *d*_R_/*d*_C_ ratio). Since radical changes are more likely to be deleterious than conservative changes, an increased *d*_R_/*d*_C_ ratio at the genome-wide scale suggests reduced selection efficiency of the population (Zuckerkandl and Pauling, 1965). A statistically significant elevation of the *d*_R_/*d*_C_ value further indicates a reduction in the effective population size (*N_e_*) of the focal lineage to the level of 10^4^-10^5^ (Zhang et al., 2024). In this case, the genome-wide *d*_R_/*d*_C_ values of flavobacteria were calculated with the software RCCalculator (Luo et al., 2017).

To implement the software, we identified single-copy orthologous genes (SCGs) shared by flavobacteria using OrthoFinder v2.2.1 (Emms and Kelly, 2015). Among the 91 flavobacterial species, we identified a total of 366 SCGs, which were then aligned using MAFFT under the auto-determined high-accuracy model. These alignments were processed in MEGA v10.2.4 (Tamura et al., 2021) to calculate the transition/transversion ratio required by RCCalculator. We then performed pairwise *d*_R_/*d*_C_ comparisons following our previous study (Luo et al., 2017). Specifically, we grouped the flavobacterial lineages into three categories: the target group (the monophyletic non-marine flavobacteria clade), the control group (the most closely related marine clade to the target group), and the reference group (the phylogenetic outgroup of the target and control groups). The *d*_R_/*d*_C_ values were calculated for comparisons between the target and reference clades, as well as between the control and reference clades. The *d*_R_/*d*_C_ values for the target and control clades were adjusted based on codon frequency and amino acid composition. Statistical tests (paired t-tests) were performed for each comparison to determine significance.

### Assessing the character dependent diversification of flavobacteria

We applied the Binary State Speciation and Extinction (BiSSE) model (Maddison et al., 2007) to estimate the diversification rate (determined by speciation and extinction rates) of marine and non-marine flavobacteria.

To reduce bias in diversification rate estimation, we expanded the flavobacteria dataset by incorporating 16S rRNA gene amplicon sequences, which allowed us to capture a broader range of uncultivated lineages. We retrieved a total of 123,523 amplicon sequences from the *Flavobacteriaceae* and *Weeksellaceae* families in the SILVA database. These sequences were then linked to NCBI nucleotide and BioSample databases to determine the habitats of the host organisms. From this, we retained 59,143 sequences with known habitats (either marine or non-marine), which were clustered into 26,700 groups using CD-HIT v4.8.1 (Fu et al., 2012) with a 0.987 identity threshold.

Next, we manually classified the habitats into marine or non-marine categories. For example, habitats like “marine,” “ocean,” “seawater,” and “gyre” were classified as marine, while those like “freshwater,” “creek,” “soil,” and “rhizosphere” were categorized as non-marine. Habitats with ambiguous descriptions, such as “water” or geographic names, were excluded from the analysis. For each cluster, we randomly selected one representative sequence, excluding those without habitat information. As a result, 17,244 representative sequences were retained, of which 3,680 sequences were longer than 1,200 bp and 13,564 sequences were shorter (Dataset S2).

We then aligned the full-length 16S rRNA sequences (>1200 bp) using MAFFT v7.453 in high accuracy mode and trimmed the alignment with trimAl v1.4 (-automated1) (Capella-Gutiérrez et al., 2009). The trimmed alignment was used for phylogenetic inference with IQ-Tree under the ‘GTR+GAMMA’ model, using the flavobacteria species tree (obtained from the section “Taxonomy sampling and phylogenomic tree reconstruction”) as the backbone. To expand the full-length 16S rRNA phylogeny, we incorporated partial 16S rRNA amplicon sequences (<1200 bp). These sequences were aligned using MAFFT with the --addfragments option and then passed to pplacer for phylogenetic inference (Matsen et al., 2010), with the alpha value for the GAMMA rate model set to 0.583 and the substitution rates for the GTR model set to 0.902, 2.527, 1.653, 0.940, 3.386, and 1.00 for AC, AG, AT, CG, CT, and GT substitutions, respectively. These parameters were estimated using IQ-Tree (Minh et al., 2020). The likelihood weight ratio (LWR) was set to 1, 0.9, and 0.8, respectively, for sensitivity testing.

To implement the BiSSE model more accurately, we approximated the posterior age estimates from the second-step dating analysis as Normal distributions using the R package “fitdistrplus” and applied them as time priors to the corresponding nodes in the 16S rRNA gene phylogeny (Table S3). This time-calibrated phylogeny was then used to reconstruct the RelTime tree using the “branch length” method in MEGA v10.2.4 (Tamura et al., 2021) (Fig. S6). Next, we used the R package *diversitree* (FitzJohn, 2012) to implement the BiSSE model with the maximum likelihood method, based on the ultrametric tree of flavobacteria. The model estimates six evolutionary parameters: speciation rates for marine and non-marine flavobacteria, extinction rates for marine and non-marine flavobacteria, and the marine-to- non-marine and non-marine-to-marine niche transition rates. These parameters were estimated under nested hypotheses: H0: all six parameters are free and differ from one another; H1: the transition rates between marine and non-marine are equal; H2: the speciation and extinction rates between marine and non-marine are equal; H3: there are no differences in the three parameters (speciation rate, extinction rate, and niche transition rate) between the two habitats (Table 1).

**Table 1.** Speciation, extinction, and habitat transition rates of flavobacteria estimated using the BiSSE model. λ, μ, and q represent the speciation rate, extinction rate, and transition rate (from the first to the second character), respectively. M and N denote marine and non-marine habitats, respectively. The four nested hypotheses are labeled as H0, H1, H2, and H3. The degrees of freedom (Df) are given in parentheses. Hypothesis marked with asterisk has the lowest AIC value at a significance level of *p* = 0.001, indicating the best BISSE model fit. The group labeled “Partial” includes only full-length 16S rRNA sequences (>1200 bp), while the “Expanded” group incorporates both full-length and partial amplicon sequences. LWR refers to the likelihood weight ratio estimated using pplacer, which reflects the robustness of sequence placement in the phylogeny. A higher LWR threshold results in fewer but more reliably placed sequences.

| Group | H (Df) | $\lambda_N$ | $\lambda_M$ | $\mu_N$ | $\mu_M$ | $q_{NM}$ | $q_{MN}$ |
| --- | --- | --- | --- | --- | --- | --- | --- |
| Expanded<br>(LWR=1) | H0 (6)* | 18.74 | 11.36 | 16.56 | 7.78 | 0.52 | 1.59 |
|  | H1 (5) | 16.64 | 12.94 | 13.38 | 10.44 | 0.96 |  |
|  | H2 (4) | 15.11 |  | 12.39 |  | 0.77 | 1.08 |
|  | H3 (3) | 15.11 |  | 12.39 |  | 0.96 |  |
| Expanded<br>(LWR=0.9) | H0 (6)* | 18.78 | 11.55 | 16.52 | 7.81 | 0.52 | 1.73 |
|  | H1 (5) | 16.57 | 13.37 | 13.13 | 10.87 | 1.01 |  |
|  | H2 (4) | 15.36 |  | 12.55 |  | 0.76 | 1.19 |
|  | H3 (3) | 15.36 |  | 12.55 |  | 1.02 |  |
| Expanded<br>(LWR=0.8) | H0 (6)* | 18.81 | 11.22 | 16.39 | 7.27 | 0.52 | 1.82 |
|  | H1 (5) | 16.50 | 13.15 | 12.78 | 10.55 | 1.03 |  |
|  | H2 (4) | 15.29 |  | 12.32 |  | 0.77 | 1.23 |
|  | H3 (3) | 15.29 |  | 12.32 |  | 1.04 |  |
| Partial<br>(>1200bp) | H0 (6)* | 12.18 | 7.31 | 9.20 | 4.53 | 0.43 | 0.74 |
|  | H1 (5) | 11.51 | 7.71 | 8.07 | 5.24 | 0.57 |  |
|  | H2 (4) | 9.84 |  | 7.21 |  | 0.54 | 0.58 |
|  | H3 (3) | 9.84 |  | 7.21 |  | 0.57 |  |

## Results

### Timetree reconstruction reveals Proterozoic habitat transitions

Our plastid-based molecular clocks (the mitochondrial-based and nuclear-based dating strategies were initially applied but later excluded in this case; see details in Methods), calibrated using 8 non-redundant eukaryotic fossil-based calibrations, 2 cyanobacterial fossil-based calibrations and 1 biomarker-based calibration (Fig. 1; Table S1), estimate the origin of flavobacteria between 2.15 Gya (posterior mean age of the total flavobacterial group; the same below) and 1.98 Gya (posterior mean age of the crown flavobacterial group; the same below), which precedes the GOE (Fig. 2a). The GOE marks a period of rising atmospheric oxygen (from <0.001% to ∼1-10% of the present atmospheric level) and enhanced organic carbon burial in marine sediments (Lyons et al., 2014). It has been proposed that low rates of organic matter burial suggest a subdued marine biosphere prior to the GOE; however, the dramatic increase in nutrient availability after the GOE likely stimulated biosphere activity (Canfield, 2021). Considering that the majority of flavobacteria are aerobic heterotrophs (García-López et al., 2019), it is plausible that the last common ancestor (LCA) of flavobacteria emerged after GOE, coinciding with the increased availability of organic carbon and nutrients in the Paleoproterozoic ocean.

**Fig. 1.**
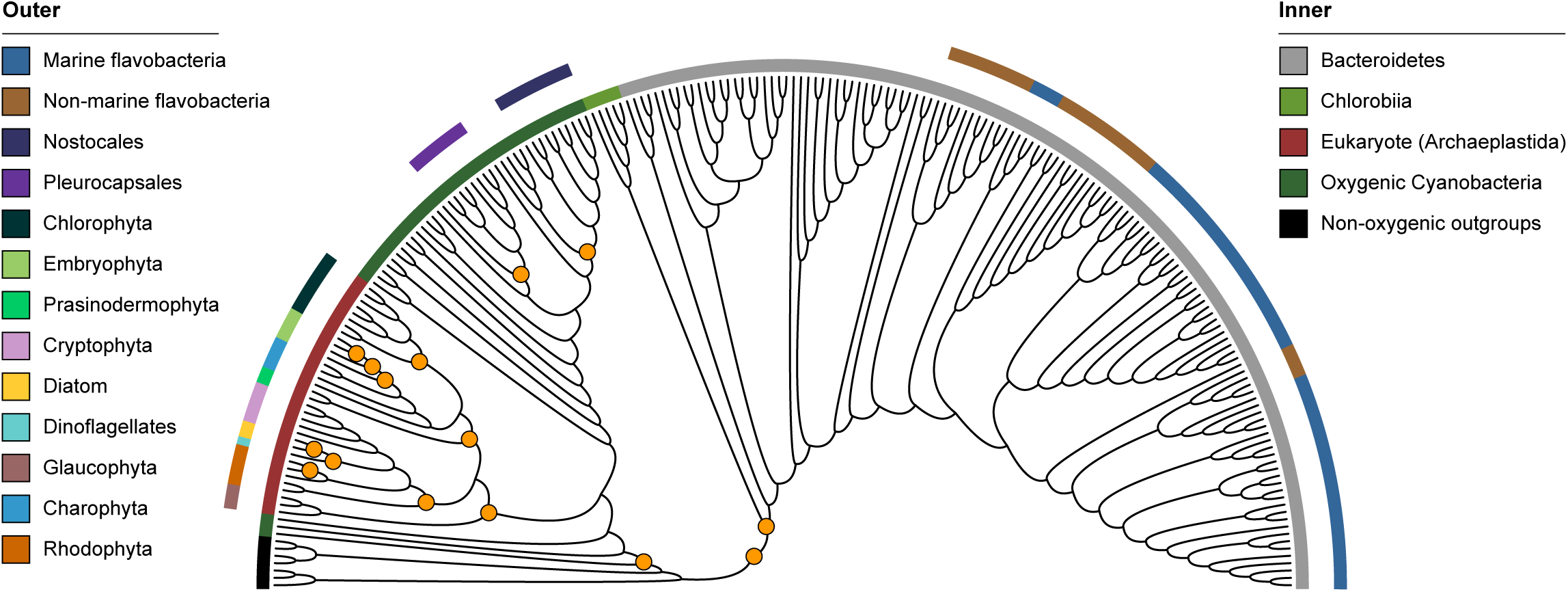
Diagram illustrating the species tree topology and the calibration points in the focal dating strategy. The inner ring represents higher-ranking taxa including different bacterial phyla and eukaryotic clade, while the outer ring denotes lower-ranking taxa, including marine/non-marine flavobacteria, and major groups within Cyanobacteria and eukaryotes. Calibrated nodes are marked with solid orange circles. Detailed calibration information is provided in Table S1.

**Fig. 2.**
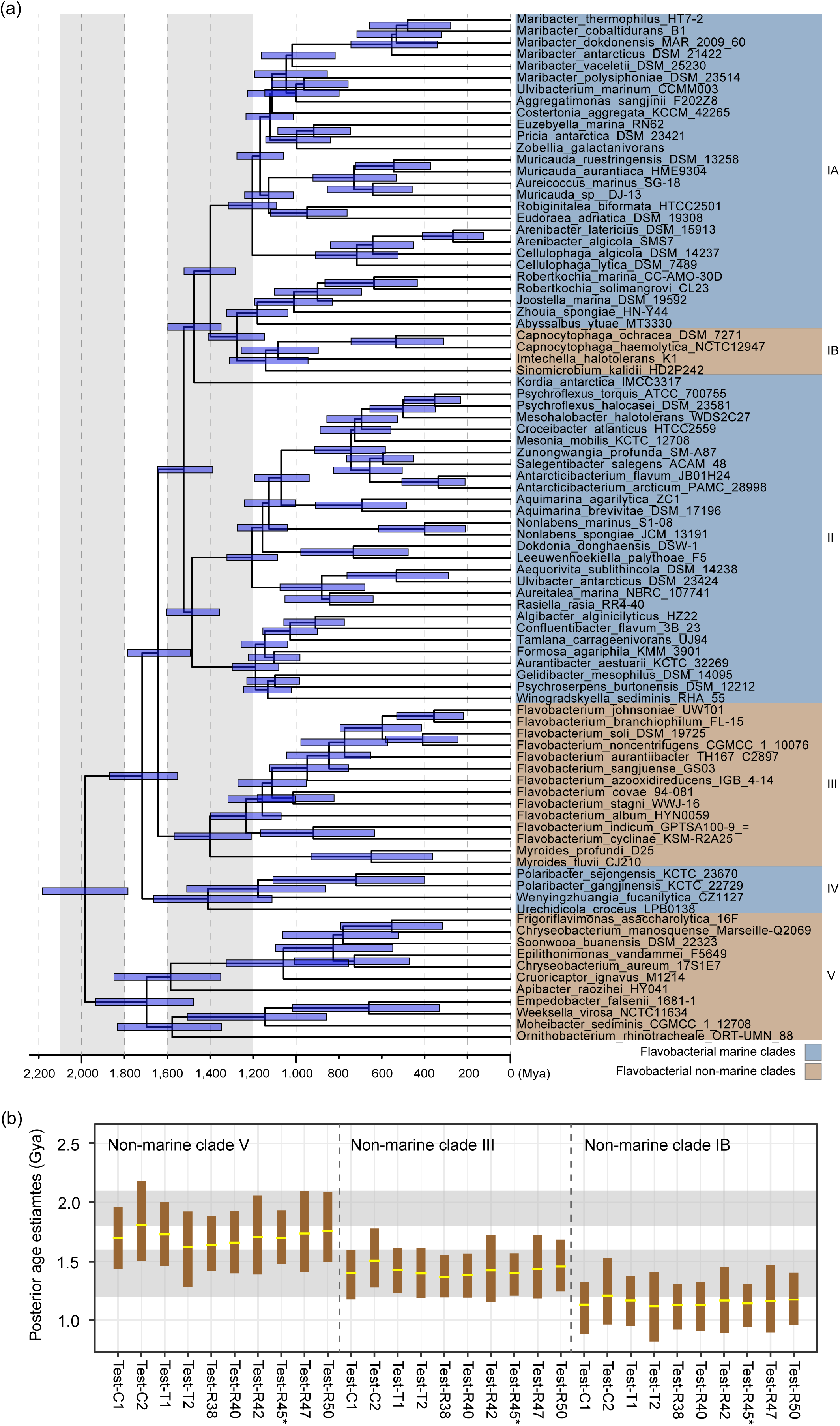
Evolutionary timeline of flavobacteria and results of sensitivity tests. (a) Chronogram of flavobacteria estimated using the focal dating strategy. Horizontal blue bars flanking ancestral nodes represent the posterior 95% highest probability density (HPD) intervals for divergence time estimates. The left and right vertical bars indicate the formation and fragmentation of the Columbia supercontinent, respectively. Marine and non-marine flavobacteria are marked in blue and brown, respectively. (b) Posterior age estimates for the last common ancestor (crown group) of non-marine clades V, III, and IB under different parameter settings. Specifically, Test-C1 and Test-C2 assess the impact of using 1.2 Gya and 2.0 Gya instead of 1.6 Gya as the minimum age constraint for the total Nostocales group. Test-T1 and Test-T2 explore alternative species tree topologies by swapping Rhodophyta and Glaucophyta, and Prasinodermophyta and Embryophyta, respectively. Test-R38, Test-R40, Test-R42, Test-R45, Test-R47, and Test-R50 examine the effect of stepwise increases in the root maximum age from 3.8 Gya to 5.0 Gya. In this panel, Test-R45 corresponds to the focal dating strategy. The upper and lower horizontal bars represent the formation and fragmentation of the Columbia supercontinent. The brown vertical bars indicate the 95% HPD intervals of posterior age estimates, with posterior means highlighted in bright yellow.

Subsequent diversification of flavobacteria involved three marine-to-non-marine transitions during the Proterozoic Eon, aligning with the life cycle of the Columbia supercontinent. Our molecular clock analysis indicates that the first marine-to-non-marine transition occurred approximately between 1.98 Gya and 1.70 Gya (Fig. 2a), spanning the formation (2.1-1.8 Gya) and expansion (1.8-1.3 Gya) of Columbia supercontinent (Zhao et al., 2002). This period was marked by major orogenic events, including the Trans-Hudson Orogeny (THO), a significant mountain-building event that contributed to the formation of the Precambrian Canadian Shield and the North American Craton at 1.83-1.80 Gya (Corrigan et al., 2009), and the Trans-North China orogeny, which led to the amalgamation of the North China Craton, crustal thickening, and uplift at ∼1.85 Gya (Liu et al., 2012). These orogenic events caused crustal shortening and thickening, progressively uplifting mountains and forming foreland basins, which later became lacustrine habitats, resembling those formed in the Andean region (Lundberg et al., 1998; Zhao et al., 2002). Therefore, we propose that the Paleo-proterozoic orogeny events created novel niches for early flavobacteria, facilitating their transition to non-marine environments.

After the assembly of the Columbia supercontinent around 1.8 Gya, it experienced prolonged subduction-driven growth through accretion along key continental margins (Zhao et al., 2004). These megamagmatic events, occurring between 1.8 and 1.7 Gya, led to the formation of distinct geological regions, including the Ketilidian Belt in Greenland and the Transscandinavian Igneous Belt in Baltica (Zhao et al., 2004). Subduction of oceanic lithosphere beneath continental plates drove mountain building both at continental margins and within intracontinental regions (Li et al., 2016), likely contributing to the formation of novel non-marine habitats for early flavobacteria.

The second and third marine-to-non-marine transitions occurred between 1.72 Gya and 1.40 Gya, as well as between 1.28 Gya and 1.14 Gya (Fig. 2a), leading to the divergence of non-marine flavobacteria clades III and IB. These transitions spanned the fragmentation of Columbia supercontinent (1.6–1.2 Gya), which were likely driven by widespread anorogenic magmatic activity (Rogers and Santosh, 2002; Zhao et al., 2004). During this period, continental rifting, characterized by crustal thinning and lithospheric stretching, occurred at divergent plate boundaries or within intracontinental settings, leading to the formation of basins and brackish transitional zones. For example, the Cryogenian Nanhua Basin formed as an intracontinental rift basin between the Yangtze and Cathaysia blocks of the South China Craton (Song et al., 2020). Recent elemental salinity proxy data indicate that the basin exhibited a systematic salinity gradient, with brackish surface waters transitioning to marine salinity at greater depths (Liu et al., 2025). Since brackish water likely represents an intermediate stage between marine and freshwater conditions, we infer that such rift-associated basins facilitated the niche transition of flavobacteria from marine to non-marine environments.

Intriguingly, this inference is supported by the phylogenomic tree of flavobacteria, as basal lineages in non-marine clades III and IB retain high salinity tolerance. In clade IB, *Sinomicrobium kalidii* and *Imtechella halotolerans* (Fig. 2a), isolated from saline-alkali environments and estuarine waters, respectively, exhibit optimal growth at ∼4% salinity (Surendra et al., 2012; Li et al., 2022). Similarly, in clade III (Fig. 2a), the basal lineage *Myroides profundi* was isolated from deep-sea sediments, with an optimal growth salinity of 1% but a tolerance of up to 8% (Zhang et al., 2008). These findings suggest that the LCA of these non-marine clades was likely adapted to a brackish environment, representing a transitional state between marine and non-marine ecotypes.

To evaluate the robustness of the flavobacterial evolutionary timeline, we conducted a series of tests under different hypotheses, focusing on key debated dating parameters. These tests included alternative calibrations for the cyanobacterial Nostocales group (marked with ‘Test-C1’ and ‘Test-C2’ in Fig. 2b) and variations in the eukaryotic species tree topology (marked with ‘Test-T1’ and ‘Test-T2’ in Fig. 2b). Additionally, to minimize potential bias introduced by the root maximum age (an obligatory parameter in the program MCMCTree), we systematically varied this constraint across a range of values: from 3.8 Gya (the earliest habitable conditions on Earth) to 4.5 Gya (Earth’s formation) and even 5.0 Gya (a methodological control without biological significance) (marked as ‘Test-R38’ to ‘Test-R50’ in Fig. 2b). These tests yielded congruent age estimates, confirming the robustness of the timeline of flavobacterial evolution. These transitions correlate with tectonic reconfigurations that fragmented marine habitats and created terrestrial niches, rather than substrate-driven competition.

### BiSSE modeling confirms asymmetric transition rates

As noted, the molecular clock-based dating analysis leverages multiple vertically transmitted orthologous genes identified from genomic sequences. Aligning the age estimates to tectonic events relies on the assumption that flavobacteria underwent repeated marine-to-non-marine habitat transitions, which was inferred based on flavobacterial species tree based on genomic sequences. To test the reliability of the transition unidirectionality while accounting for the uncultivated majority that lacks genomic sequences, we applied the Binary-State Speciation and Extinction (BiSSE) model to time-calibrated 16S rRNA phylogenies. In the phylogeny based on full-length 16S rRNA sequences (3,680 in total), we observed an interleaved distribution of marine and non-marine flavobacterial clades, consistent with the pattern found in the phylogeny based on concatenated ribosomal proteins (Zhang et al., 2019) (Fig. S6a). This distribution remained stable even when the dataset was expanded to include many more additional 16S amplicon sequences (13,564 in total) (Fig. S6b). Based on the expanded dataset, we found that non-marine lineages exhibited higher turnover (speciation + extinction rates: 35.3 vs. 19.14 events/Gya) but lower net diversification (speciation - extinction rates: 2.18 vs. 3.58) compared to their marine counterparts (Table 1; ‘Expanded’; Likelihood Weight Ratio or LWR =1), reflecting the instability of fragmented non-marine habitats. For example, soil flavobacteria face stochastic moisture and nutrient availability, driving boom-bust population dynamics (Fierer et al., 2007). Similarly, temperatures are more heterogeneous across land surfaces than across ocean surfaces (Burrows et al., 2011), imposing strong selective pressures that drive the speciation of non-marine flavobacteria. Finally, anthropogenic barriers, such as agricultural and urbanized areas, are primarily located in terrestrial habitats, hindering migration and facilitating speciation (Mendenhall et al., 2012; Knapp et al., 2017).

Critically, transition rates were profoundly asymmetric: marine-to-non-marine shifts (1.59 events/Gya) outpaced reverse transitions (0.52 events/Gya; *p* < 0.001). This unidirectionality suggests that once flavobacteria colonized non-marine habitats, reverse transitions became evolutionarily constrained. Following habitat transitions, the effective population sizes of the transitioned species may be reduced due to population bottlenecks and/or the founder effect, and gene flow becomes restricted due to the fragmented nature of many non-marine habitats. These processes are known to reduce the efficacy of purifying selection acting to eliminate deleterious mutations, thereby allowing the accumulation of mildly deleterious mutations and resulting in the ‘ratchet effect’ (Moran, 1996). Here, we used the genome-averaged *d_R_*/*d_C_* ratio (the ratio of radical to conservative nonsynonymous substitution rates) as a proxy for selection efficiency, which was validated in our prior studies that demonstrated genetic drift as a major driver of *Prochlorococcus* genome reduction (Luo et al., 2017; Zhang et al., 2024). We show that an elevated genome-averaged *d_R_*/*d_C_* ratio is associated with the non-marine clade IB (Fig. 3). This is perhaps not surprising, given that most clade IB flavobacteria (*Capnocytophaga*) are associated with eukaryotic hosts (Butler, 2015). In light of the observation that non-marine host-associated flavobacteria generally undergo genome reduction (Zhang et al., 2019; Lee et al., 2023), our analysis suggests that genome-wide evolutionary changes make transitions from non-marine to marine habitats challenging, following marine-to-non-marine transitions and new habitat adaptation. This process mirrors genome erosion in obligate symbionts (Moran, 1996) and helps explain the rarity of non-marine-to-marine reversals.

**Fig. 3.**
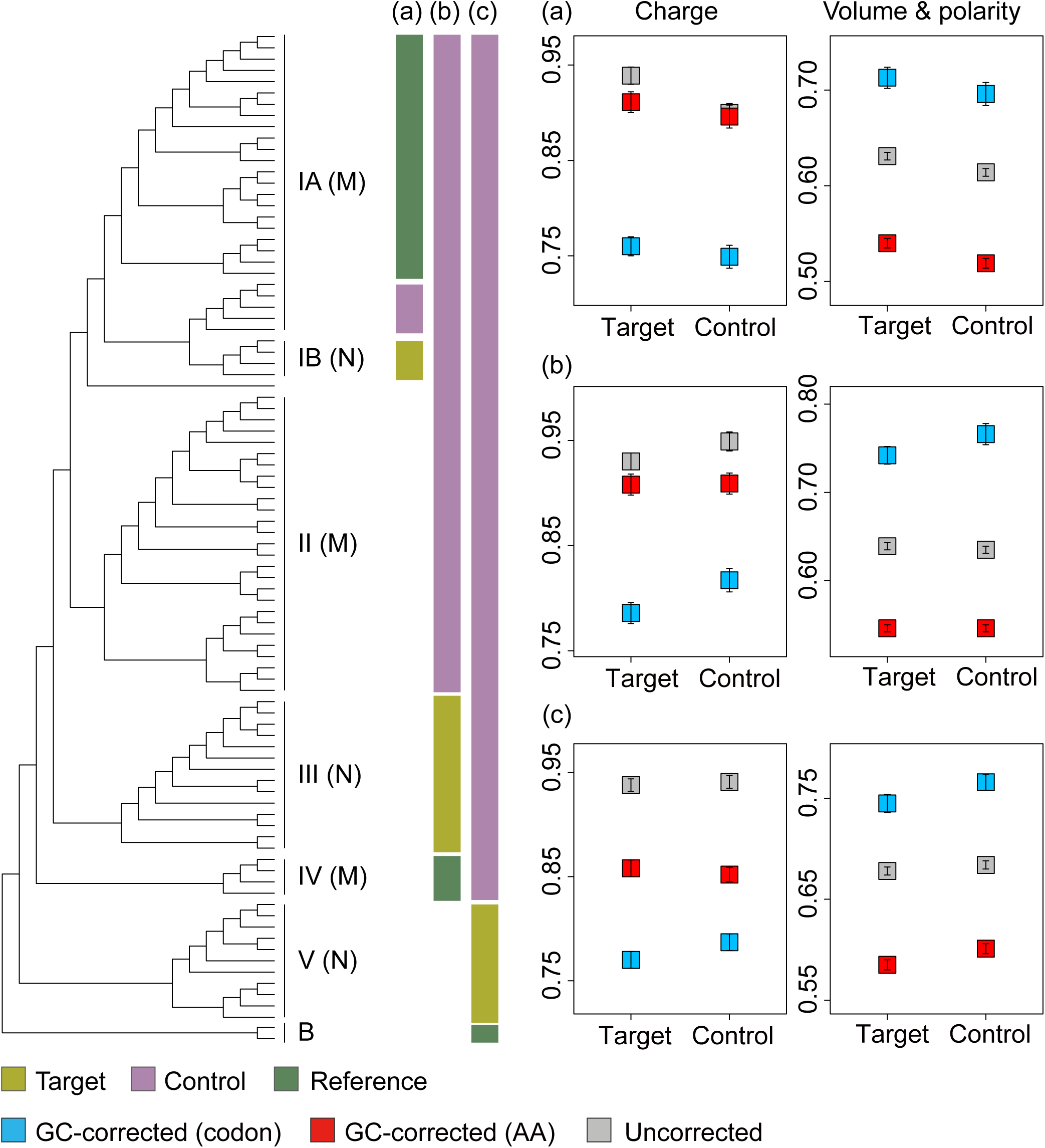
Diagram illustrating the schemes and the results of *d*_R_/*d*_C_ comparison. The genome-wide means of *d*_R_/*d*_C_ values at the ancestral branches (‘Target’) leading to (a) non-marine clade IB, (b) non-marine clade III, and (c) non-marine clade V compared with that at the sister lineages (‘Control’). The *d*_R_/*d*_C_ values were classified based on the physicochemical classification of the amino acids by charge or by volume and polarity, and were either GC-corrected by codon frequency (blue), GC-corrected by amino acid (AA) frequency (red) or uncorrected (gray). Error bars of *d*_R_/*d*_C_ represent the standard error of the mean.

Notably, we conducted sensitivity tests on the input dataset for the BiSSE model. Using the likelihood weight ratio (LWR) as a threshold to assess the robustness of amplicon sequence placement on the phylogeny, we compiled three datasets that included both full-length 16S rRNA genes and amplicon sequences (denoted as ‘Expanded’ in Table 1). Besides, we performed another test using only full-length 16S rRNA gene sequences as input data for the model (denoted as ‘Partial’ in Table 1). While there were differences in exact speciation and extinction rates, all tests supported the above findings, reinforcing our conclusion regarding the higher turnover rate and lower net diversification rate of non-marine flavobacteria, as well as the genome erosion hypothesis related to the marine-to-non-marine niche transition.

## Discussion

### Reconciling tectonics and microbial evolution

Our findings redefine the ecological trajectory of flavobacteria, positioning Proterozoic tectonics, rather than substrate-specific enzyme innovation, as the catalyst for their evolutionary success. The post-GOE origin of flavobacteria suggests that they may have utilized the organic carbon released by Cyanobacteria in their early stages. The subsequent diversification of non-marine flavobacteria was likely driven by the tectonic reconfiguration of Earth’s landscapes, which was likely unrelated to glycans. In oceans, phytoplankton and macroalgae form the foundation of the food web and serve as the primary sources of polysaccharides (Arnosti et al., 2021). In both groups, polysaccharides are key structural and storage compounds, accounting for over 50% of their dry mass, although their composition varies between taxa (Mabeau and Kloareg, 1987; Becker et al., 2020). Additionally, exopolysaccharides produced by mucus and biofilms from various marine organisms contribute to the diversity and abundance of polysaccharides in the ocean (Arnosti et al., 2021). In non-marine niches, polysaccharides constitute about a quarter of soil organic matter, much of which originates from plant roots and debris (Oades, 1984). However, with the exception of Cyanobacteria, the major contributors of the organic carbon have not been evolved at that time, such as the phytoplanktonic diatom (Becker et al., 2020) and the macroalgae species of *Florideophyceae* (Hill et al., 2015), strengthening our hypothesis that extrinsic geological factors, rather than glycan specialization, drove the early evolution and marine-to-non-marine niche transition of flavobacteria.

Strikingly, the scenario that supercontinental cycles generated physical barriers (e.g., mountain ranges) and novel habitats (e.g., rift basins) for speciation parallels that on eukaryotic macroevolution. For example, the breakup of Pangaea was considered to drove the diversification of angiosperms and mammals by creating biogeographic provinces (Chaboureau et al., 2014; Jordan et al., 2016). Our results extend this paradigm to microbes, emphasizing that planetary-scale geology shapes not only macroscopic biodiversity but also microbial metabolic networks.

### Implications for carbon cycling

As major mineralizers of organic matter, the increased abundance of flavobacteria has been linked to enhanced primary production (Williams et al., 2013). Before GOE, only 5-10% of organic matter was buried in marine sediments compared to today. In contrast, after the GOE, there was a dramatic increase in nutrient availability, which stimulated biosphere activity (Canfield, 2021). The GOE’s organic carbon burial created a reservoir of recalcitrant DOC, which flavobacteria’s CAZymes unlocked, releasing CO_2_ and nutrients back into ecosystems. Therefore, flavobacteria’s tectonically driven diversification had profound implications for Proterozoic carbon cycling.

The later terrestrialization of flavobacteria coincided with the expansion of terrestrial ecosystems, during which flavobacteria may have accelerated soil organic matter turnover, contributing to carbon cycling in early terrestrial habitats. Modern observations provide further evidence. For example, *Flavobacterium ginsengisoli* displayed an extremely high degradation capacity for lignocellulose, which accounts for 20-50% of agricultural waste (Van Dyk and Pletschke, 2012). Thus, their Proterozoic radiation likely established feedbacks between tectonics, microbial evolution, and climate that persist today.

### Broader lessons for microbial biogeography

Our study challenges the prevailing view that microbial biogeography is shaped primarily by environmental selection (De Wit and Bouvier, 2006; Milke et al., 2022). While niche-specific traits mediate local adaptation, flavobacteria’s global distribution reflects ancient tectonic processes that created the niches themselves. This “top-down” control (from geological events to habitat availability and then to microbial divergence) complements “bottom-up” drivers (e.g., pH, temperature) and underscores the need to integrate deep-time perspectives into microbial ecology. For example, the Andean uplift (∼20 Mya) created altitudinal gradients that structured *Pseudomonas* diversity (Chauhan et al., 2023). In the same way, Proterozoic orogenies may have generated beta diversity hotspots for flavobacteria.

## Conclusion

By integrating fossil-calibrated molecular clocks with genomic and biogeographic data, we demonstrate that flavobacteria’s deep divergence was tectonically catalyzed, not substrate-driven. Their Proterozoic diversification underscores the profound influence of geological dynamics on microbial evolution, offering a framework to predict how contemporary environmental shifts may reshape microbial contributions to global carbon cycling. As anthropogenic activities alter landscapes at tectonic scales, understanding these microbe-geology feedbacks will be critical for managing ecosystem resilience in a changing climate.

## Data availability

The datasets generated during the current study are available in the GitHub repository (https://github.com/luolab-cuhk/Flavo-dating).

## Acknowledgements

We would like to thank Dr. Sishuo Wang and Dr. Tianhua Liao for their assistance with the molecular clock analysis. This work is supported by the Hong Kong Research Grants Council (RGC) General Research Fund (GRF) (14110820), and the Natural Science Foundation of Guangdong Province, China (2022A1515010844 to HZ).

## Notes

https://github.com/luolab-cuhk/Flavo-dating

### Competing Interest Statement

The authors have declared no competing interest.

