## Supplementary material for "Proterozoic Tectonic Drivers Underpin Flavobacterial Diversification": SI

*Corresponding author: Haiwei Luo

**This PDF file includes:**

Figure S1-S6

Table S1-S3

Fig. S1

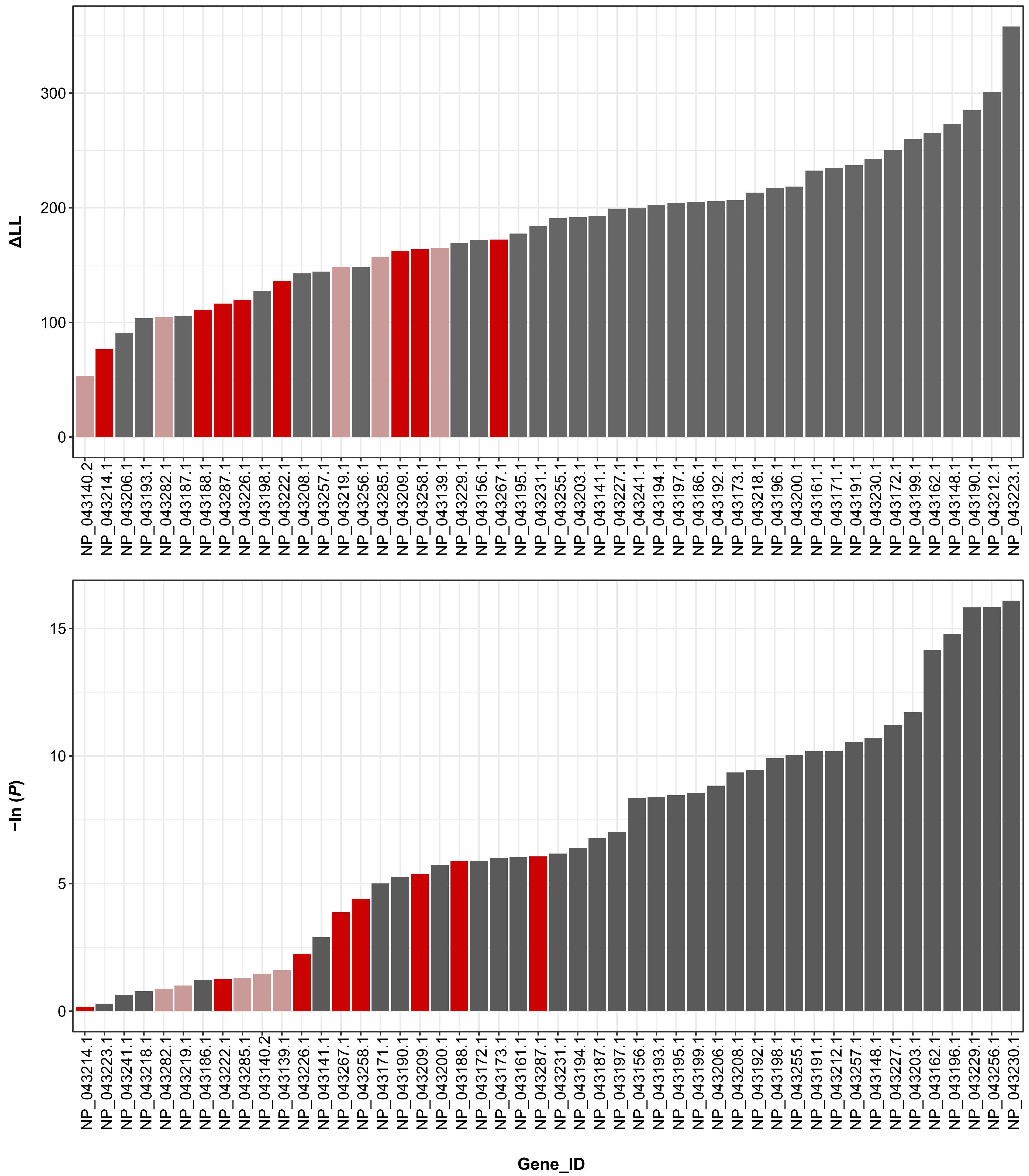

Fig. S1 Rankings of (a) gene verticality, using ΔLL as a proxy, and (b) evolutionary rate differences between bacterial and eukaryotic groups, using the log-transformed p-value of the Wilcoxon signed-rank test as a proxy. Gene families marked in red and pink appear in the first half of both rankings, but pink-marked genes were excluded from dating analysis due to the large phylogenetic incongruence between gene tree and the species tree.

Fig. S2

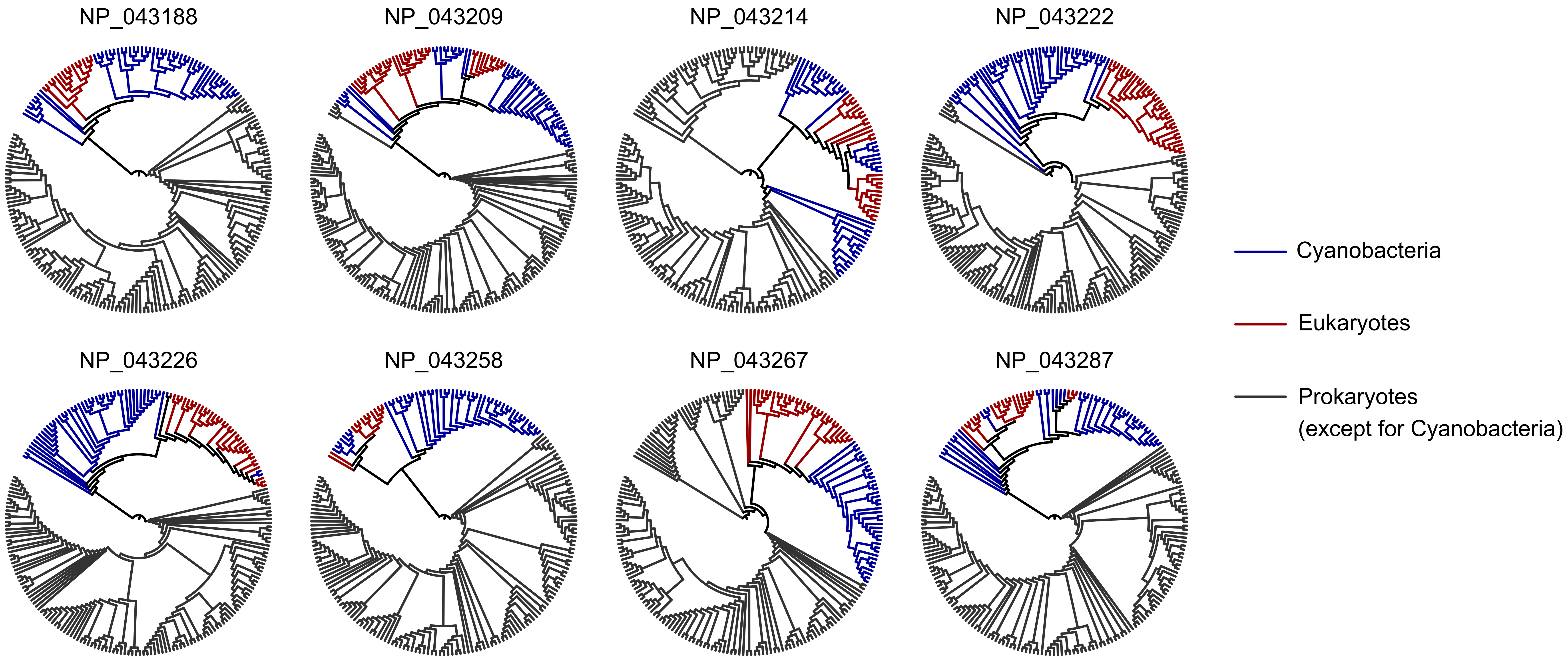

Fig. S2 The phylogeny of gene families exhibiting high gene verticality and low differences in evolutionary rate between bacterial and eukaryotic groups. Genes suspected to be derived from horizontal gene transfer were manually checked and removed. Cyanobacterial and eukaryotic species are marked in blue and red, respectively, while other bacterial species are shown in black.

Fig. S3

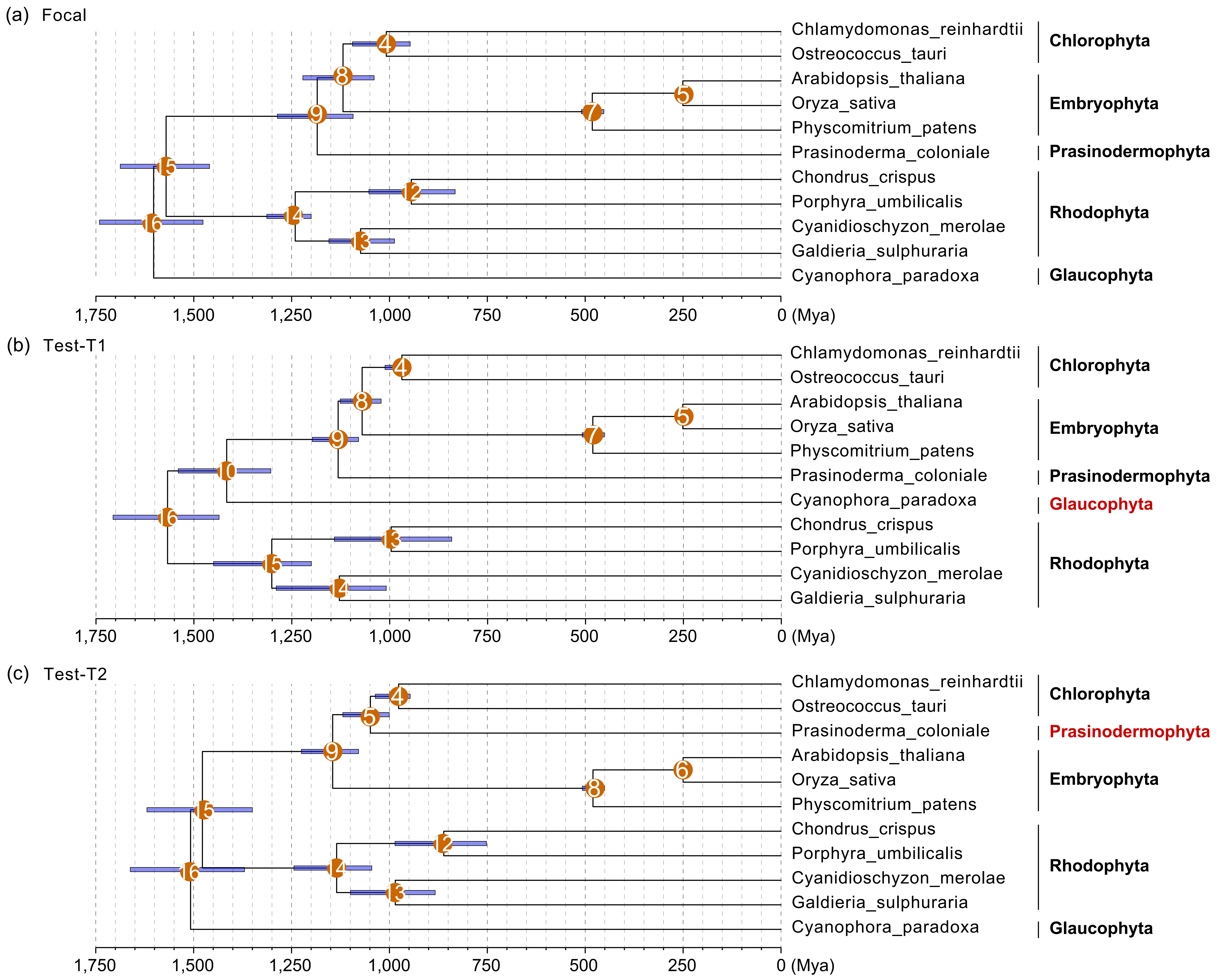

Fig. S3 The posterior age estimates of eukaryotic species derived from the first-step dating analysis, which was based on the eukaryotic portion of the species tree and eukaryotic fossil-based calibrations. The data were obtained from our recent study (Zhang et al., 2023). Panels (a), (b), and (c) depict the focal species tree and the alternative species trees ‘Test-T1’ and ‘Test-T2,’ respectively. Eukaryotic groups with differing phylogenetic positions between the focal and alternative species trees are highlighted in red. Solid orange circles indicate calibrated nodes, with numbers corresponding to Table S1.

Fig. S4 The posterior age estimates of both bacterial and eukaryotic species derived from the second-step dating analysis, which incorporated both eukaryotic and bacterial groups using eukaryotic fossil-based calibrations, cyanobacterial fossil-based calibrations, and biomarker-based calibration. The flanking horizontal blue bars on ancestral nodes represent the posterior 95% highest probability density (HPD) interval of the estimated divergence time. Solid orange circles denote calibrated nodes, with numbers corresponding to Table S1. Panels (a), (b), and (c) correspond to the focal dating strategy, alternative species tree topology 'Test-T1,' and alternative species tree topology 'Test-T2,' respectively.

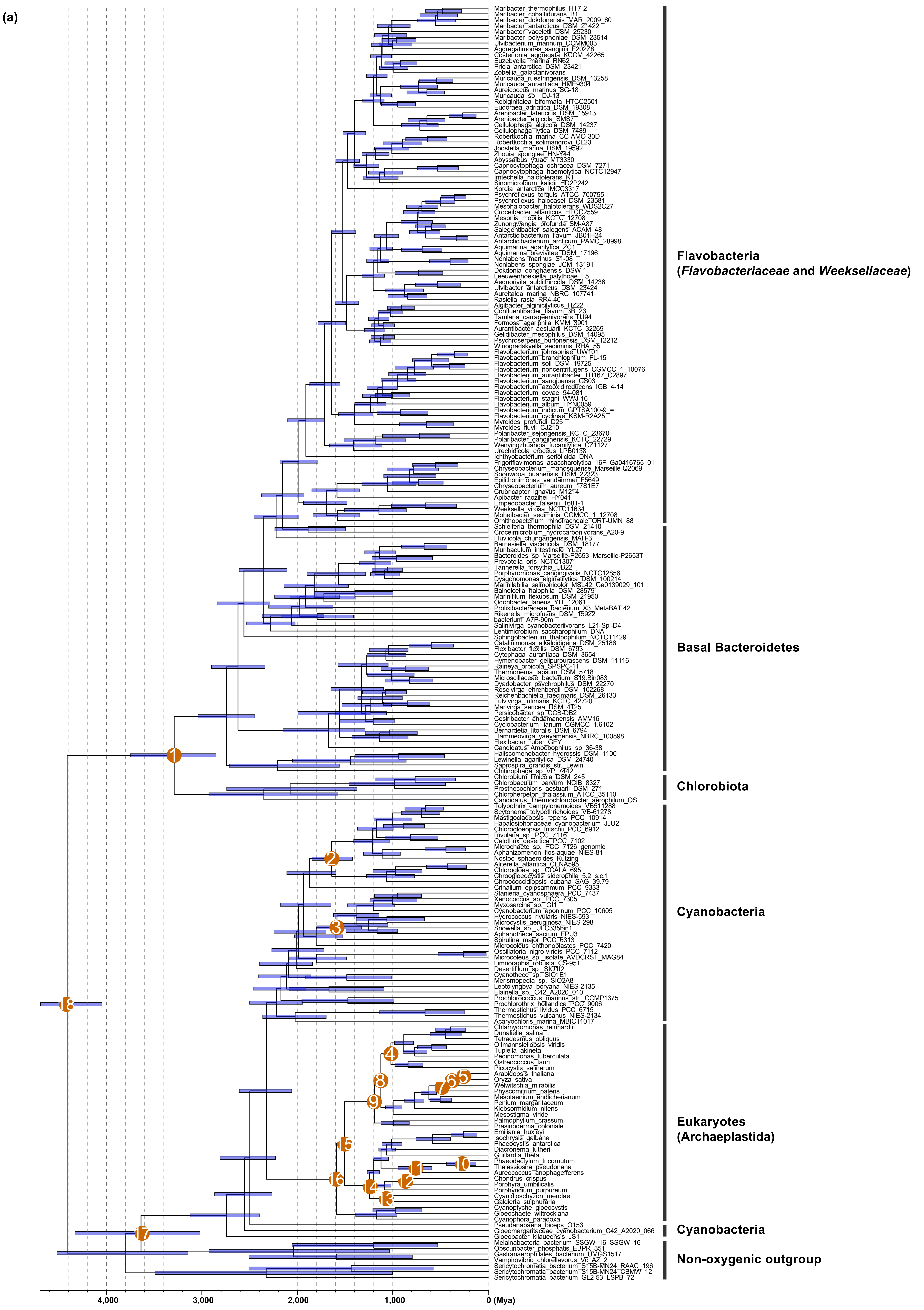

(b)

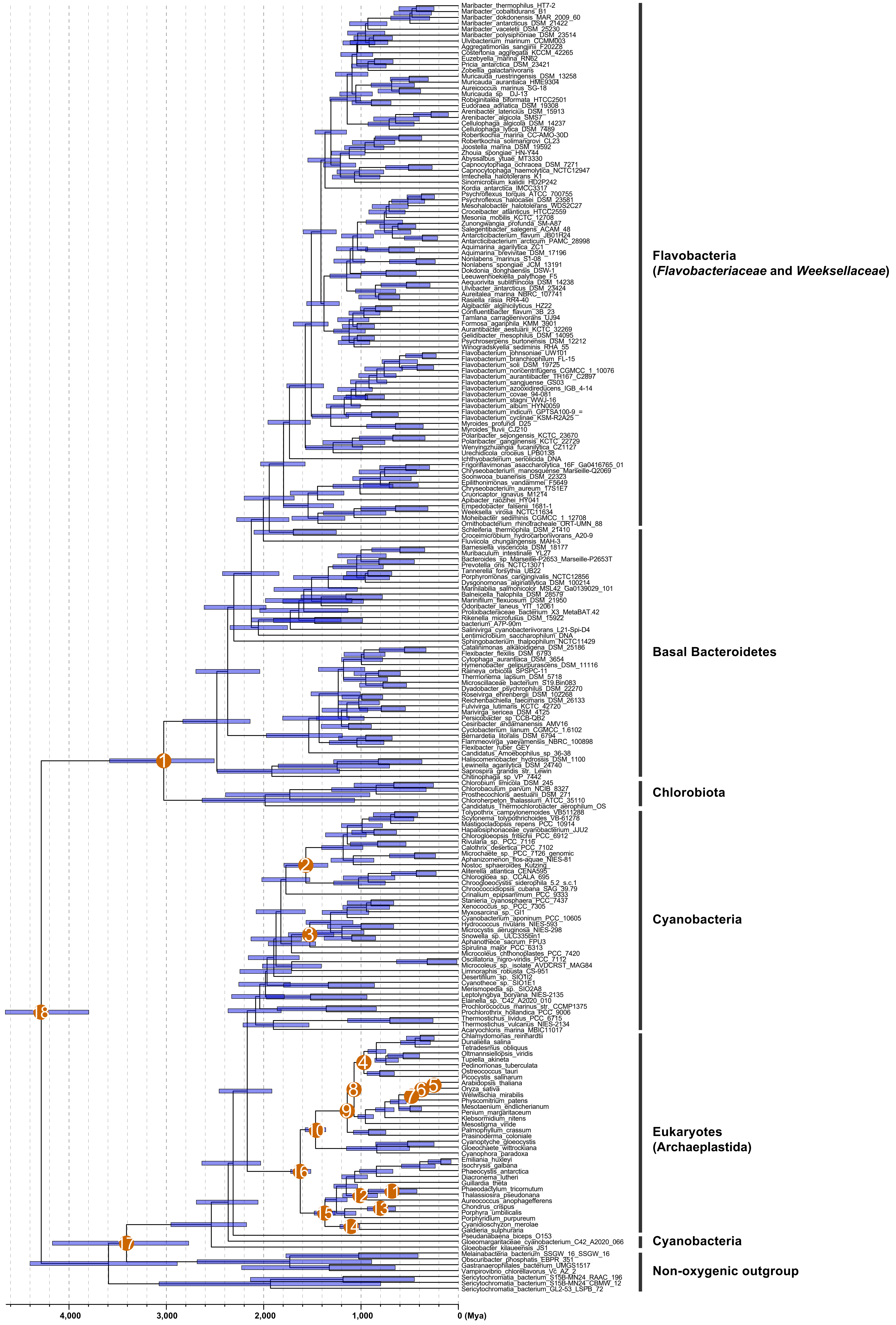

Maribacter_thermophilus_HT7-2
Maribacter_cobaltidurans_B1
Maribacter_dokdonensis_MAR_2009_60
Maribacter_antarcticus_DSM_21422
Maribacter_vaceletii_DSM_25230
Maribacter_polysiphoniae_DSM_23514
Ulvibacterium_marinum_CCMM003
Aggregatimonas_sangjinii_F202Z8
Costertonia_aggregata_KCCM_42265
Euzebyella_marina_RN62
Pricia_antarctica_DSM_23421
Zobellia_galactanivorans
Muricauda_ruestringensis_DSM_13258
Muricauda_aurantiaca_HME9304
Aureicoccus_marinus_SG-18
Muricauda_sp._DJ-13
Robiginitalea_biformata_HTCC2501
Eudoraea_adriatica_DSM_19308
Arenibacter_latericius_DSM_15913
Arenibacter_algicola_SMS7
Cellulophaga_algicola_DSM_14237
Cellulophaga_lytica_DSM_7489
Robertkochia_marina_CC-AMO-30D
Robertkochia_solimangrovi_CL23
Joostella_marina_DSM_19592
Zhouia_spongiae_HN-Y44
Abyssalbus_ytuae_MT3330
Capnocytophaga_ochracea_DSM_7271
Capnocytophaga_haemolytica_NCTC12947
Imtechella_halotolerans_K1
Sinomicrobium_kalidii_HD2P242
Kordia_antarctica_IMCC3317
Psychroflexus_torquis_ATCC_700755
Psychroflexus_halocasei_DSM_23581
Mesohalobacter_halotolerans_WDS2C27
Croceibacter_atlanticus_HTCC2559
Mesonia_mobilis_KCTC_12708
Zunongwangia_profunda_SM-A87
Salegentibacter_salegens_ACAM_48
Antarcticibacterium_flavum_JB01R24
Antarcticibacterium_arcticum_PAMC_28998
Aquimarina_agarilytica_ZC1
Aquimarina_brevivitae_DSM_17196
Nonlabens_marinus_S1-08
Nonlabens_spongiae_JCM_13191
Dokdonia_donghaensis_DSW-1
Leeuwenhoekiella_palythoae_F5
Aequorivita_sublithincola_DSM_14238
Ulvibacter_antarcticus_DSM_23424
Aureitalea_marina_NBRC_107741
Rasiella_rasia_RR4-40
Algibacter_alginicilyticus_HZ22
Confluentibacter_flavum_3B_23
Tamlana_carrageenivorans_UJ94
Formosa_agariphila_KMM_3901
Aurantibacter_aestuarii_KCTC_32269
Gelidibacter_mesophilus_DSM_14095
Psychroserpens_burtonensis_DSM_12212
Winogradskyella_sediminis_RHA_55
Flavobacterium_johnsoniae_UW101
Flavobacterium_branchiophilum_FL-15
Flavobacterium_soli_DSM_19725
Flavobacterium_noncentrifugens_CGMCC_1_10076
Flavobacterium_aurantiibacter_TH167_C2897
Flavobacterium_sangjuense_GS03
Flavobacterium_azooxidireducens_IGB_4-14
Flavobacterium_covae_94-081
Flavobacterium_stagni_WWJ-16
Flavobacterium_album_HYN0059
Flavobacterium_indicum_GPTSA100-9_=
Flavobacterium_cyclinae_KSM-R2A25
Myroides_profundi_D25
Myroides_fluvii_CJ210
Polaribacter_sejongensis_KCTC_23670
Polaribacter_gangjinensis_KCTC_22729
Wenyingzhuangia_fucanilytica_CZ1127
Urechidicola_croceus_LPB0138
Ichthyobacterium_seriolicida_DNA
Frigoriflavimonas_asaccharolytica_16F_Ga0416765_01
Chryseobacterium_manosquense_Marseille-Q2069
Soonwooa_buanensis_DSM_22323
Epilithonimonas_vandammei_F5649
Chryseobacterium_aureum_17S1E7
Cruoricaptor_ignavus_M1214
Apibacter_raozihei_HY041
Empedobacter_falsenii_1681-1
Weeksella_virosa_NCTC11634
Moheibacter_sediminis_CGMCC_1_12708
Ornithobacterium_rhinotracheale_ORT-UMN_88
Schleiferia_thermophila_DSM_21410
Croceimicrobium_hydrocarbonivorans_A20-9
Fluviicola_chungangensis_MAH-3
Barnesiella_viscericola_DSM_18177
Muribaculum_intestinale_YL27
Bacteroides_sp_Marseille-P2653_Marseille-P2653T
Prevotella_oris_NCTC13071
Tannerella_forsythia_UB22
Porphyromonas_cangingivalis_NCTC12856
Dysgonomonas_alginatilytica_DSM_100214
Marinilabilia_salmonicolor_MSL42_Ga0139029_101
Balneicella_halophila_DSM_28579
Marinifilum_flexuosum_DSM_21950
Odoribacter_laneus_YIT_12061
Prolixibacteraceae_bacterium_X3_MetaBAT.42
Rikenella_microfusus_DSM_15922
bacterium_A7P-90m
Salinivirga_cyanobacteriivorans_L21-Spi-D4
Lentimicrobium_saccharophilum_DNA
Sphingobacterium_thalpophilum_NCTC11429
Catalinimonas_alkaloidigena_DSM_25186
Flexibacter_flexilis_DSM_6793
Cytophaga_aurantiaca_DSM_3654
Hymenobacter_gelipurpurascens_DSM_11116
Raineya_orbicola_SPSPC-11
Thermonema_lapsum_DSM_5718
Microscillaceae_bacterium_S19.Bin083
Dyadobacter_psychrophilus_DSM_22270
Roseivirga_ehrenbergii_DSM_102268
Reichenbachiella_faecimaris_DSM_26133
Fulvivirga_lutimaris_KCTC_42720
Marivirga_sericea_DSM_4125
Persicobacter_sp_CCB-QB2
Cesiribacter_andamanensis_AMV16
Cyclobacterium_lianum_CGMCC_1.6102
Bernardetia_litoralis_DSM_6794
Flammeovirga_yaeyamensis_NBRC_100898
Flexibacter_ruber_GEY
Candidatus_Amoebophilus_sp_36-38
Haliscomenobacter_hydrossis_DSM_1100
Lewinella_agarilytica_DSM_24740
Saprospira_grandis_str._Lewin
Chitinophaga_sp_VP_7442
Chlorobium_limicola_DSM_245
Chlorobaculum_parvum_NCIB_8327
Prosthecochloris_aestuarii_DSM_271
Chloroherpeton_thalassium_ATCC_35110
Candidatus_Thermochlorobacter_aerophilum_OS
Tolypothrix_campylonemoides_VB511288
Scytonema_tolypothrichoides_VB-61278
Mastigocladopsis_repens_PCC_10914
Hapalosiphonaceae_cyanobacterium_JJU2
Chlorogloeopsis_fritschii_PCC_6912
Rivularia_sp._PCC_7116
Calothrix_desertica_PCC_7102
Microchaete_sp._PCC_7126_genomic
Aphanizomenon_flos-aquae_NIES-81
Nostoc_sphaeroides_Kutzing
Aliterella_atlantica_CENA595
Chroogloea_sp._CCALA_695
Chroogloeocystis_siderophila_5.2_s.c.1
Chroococcidiopsis_cubana_SAG_39.79
Crinalium_epipsammum_PCC_9333
Stanieria_cyanosphaera_PCC_7437
Xenococcus_sp._PCC_7305
Myxosarcina_sp._GI1
Cyanobacterium_aponinum_PCC_10605
Hydrococcus_rivularis_NIES-593
Microcystis_aeruginosa_NIES-298
Snowella_sp._ULC335bin1
Aphanothece_sacrum_FPU3
Spirulina_major_PCC_6313
Microcoleus_chthonoplastes_PCC_7420
Oscillatoria_nigro-viridis_PCC_7112
Microcoleus_sp._isolate_AVDCRST_MAG84
Limnoraphis_robusta_CS-951
Desertifilum_sp._SIOT2
Cyanothece_sp._SIO1E1
Merismopedia_sp._SIO2A8
Leptolyngbya_boryana_NIES-2135
Elainella_sp._C42_A2020_010
Prochlorococcus_marinus_str._CCMP1375
Prochlorothrix_hollandica_PCC_9006
Thermostichus_lividus_PCC_6715
Thermostichus_vulcanus_NIES-2134
Acaryochloris_marina_MBIC11017
Chlamydomonas_reinhardtii
Dunaliella_salina
Tetradesmus_obliquus
Oltmannsiellopsis_viridis
Tupiella_akineta
Pedinomonas_tuberculata
Ostreococcus_tauri
Picocystis_salinarum
Arabidopsis_thaliana
Oryza_sativa
Welwitschia_mirabilis
Physcomitrium_patens
Mesotaenium_endlicherianum
Penium_margaritaceum
Klebsormidium_nitens
Mesostigma_viride
Palmophyllum_crassum
Prasinoderma_coloniale
Cyanoptyche_gloeocystis
Gloeochaete_wittrockiana
Cyanophora_paradoxa
Emiliania_huxleyi
Isochrysis_galbana
Phaeocystis_antarctica
Diacronema_lutheri
Guillardia_theta
Phaeodactylum_tricornutum
Thalassiosira_pseudonana
Aureococcus_anophagefferens
Chondrus_crispus
Porphyra_umbilicalis
Porphyridium_purpureum
Cyanidioschyzon_merolae
Galdieria_sulphuraria
Pseudanabaena_biceps_O153
Gloeomargaritaceae_cyanobacterium_C42_A2020_066
Gloeobacter_kilaueensis_JS1
Melainabacteria_bacterium_SSGW_16_SSGW_16
Obscuribacter_phosphatis_EBPR_351
Gastranaerophilales_bacterium_UMGS1517
Vampirovibrio_chlorellavorus_Vc_AZ_2
Sericytochromatia_bacterium_S15B-MN24_RAAC_196
Sericytochromatia_bacterium_S15B-MN24_CBMW_12
Sericytochromatia_bacterium_GL2-53_LSPB_72
Flavobacteria
(Flavobacteriaceae and Weeksellaceae)
Basal Bacteroidetes
Chlorobiota
Cyanobacteria
Eukaryotes
(Archaeplastida)
Cyanobacteria
Non-oxygenic outgroup
4,000
3,000
2,000
1,000
0 (Mya)

(c)

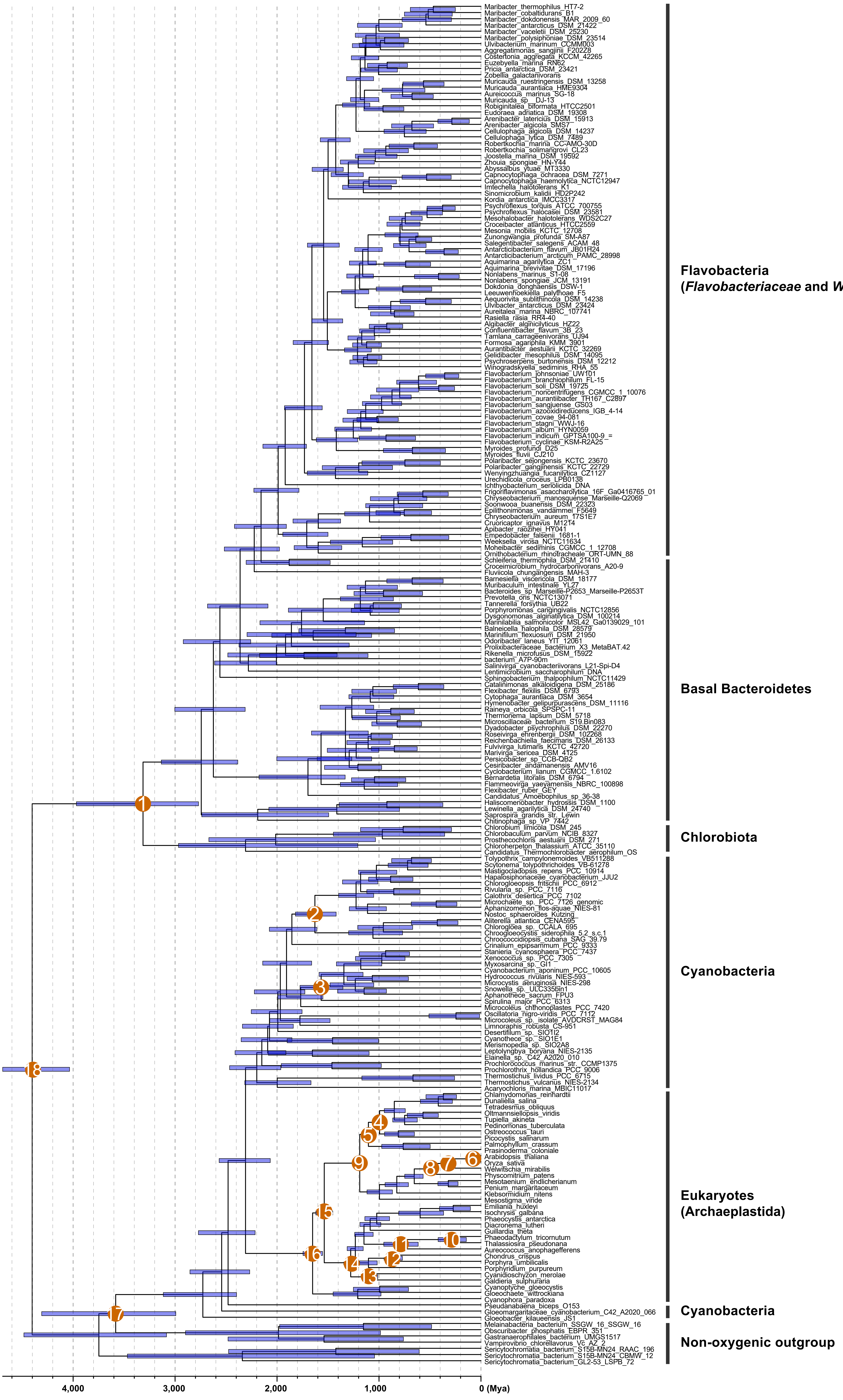
Flavobacteria
(*Flavobacteriaceae* and *Weeksellaceae*)
Basal Bacteroidetes
Chlorobiota
Cyanobacteria
Eukaryotes
(Archaeplastida)
Cyanobacteria
Non-oxygenic outgroup
4,000
3,000
2,000
1,000
0 (Mya)

Fig. S5

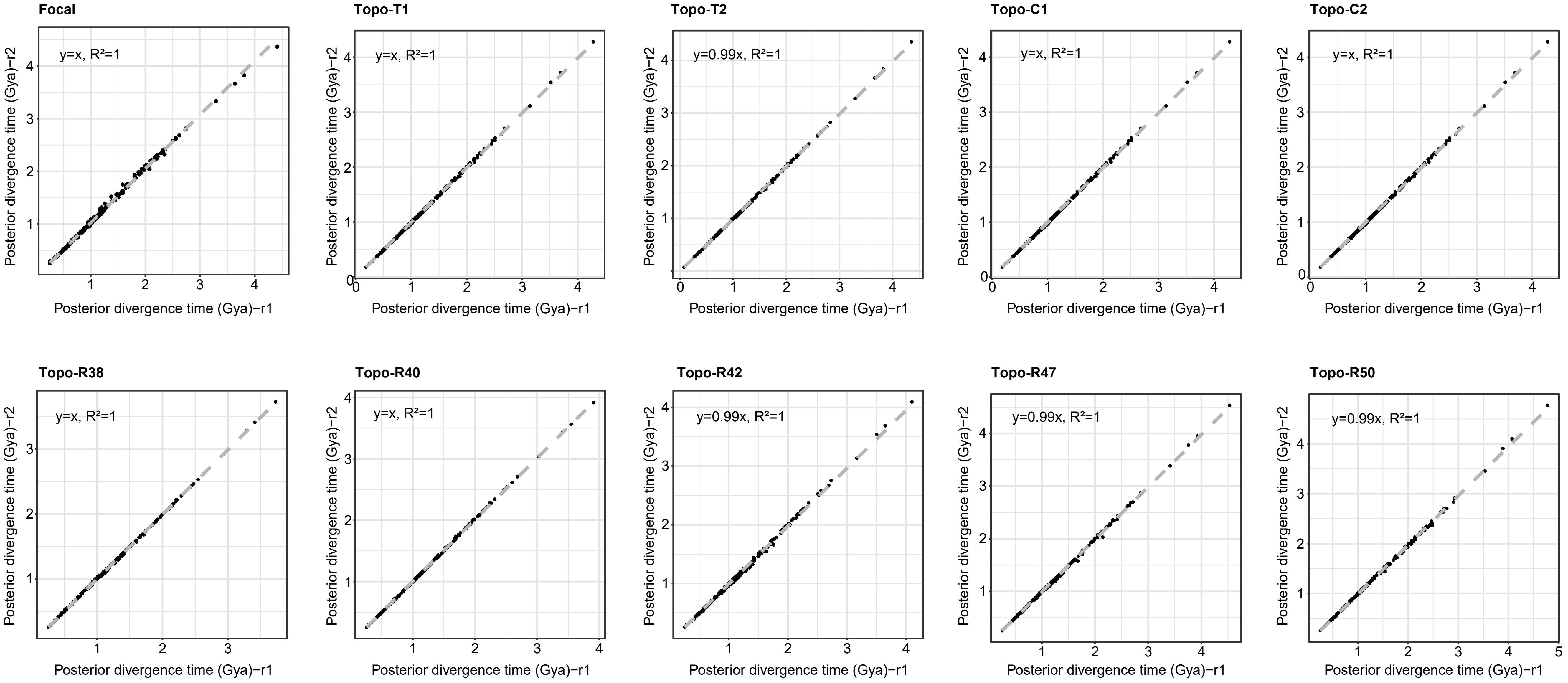

Fig. S5 The diagnostics plots showing the convergence of the MCMC analyses. Each dating analysis was run twice to confirm convergence. The dots in the plots represent ancestral nodes in the species tree, and their positions along the y = x line indicate a high level of convergence. The labels ‘Focal,’ ‘Topo-T1,’ ‘Topo-T2,’ ‘Topo-C1,’ ‘Topo-C2,’ and ‘Topo-R38’ to ‘Topo-R50’ correspond to those shown in Fig. 2b.

Fig. S6

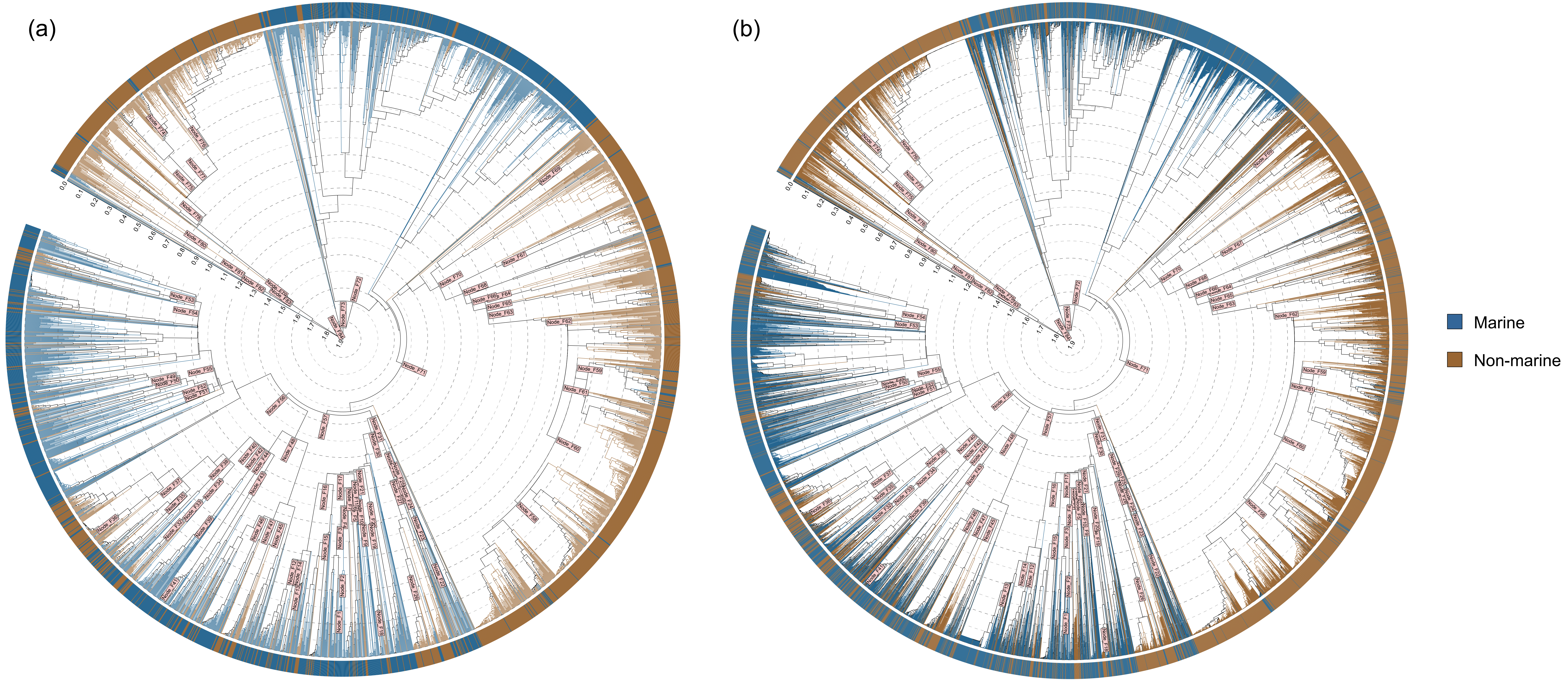

Fig. S6 The chronogram of flavobacteria (16S rRNA gene-based) was inferred using the RelTime method implemented in MEGA. The posterior ages derived from the focal dating analysis of flavobacteria (genome-based; Fig. S4a) were approximated to normal distributions (Table S3), which were then used as time priors in the RelTime timetree reconstruction. Divergence time estimation was based on (a) 3,680 full-length 16S rRNA genes and (b) 13,564 full-length and partial amplicon 16S rRNA sequences. Flavobacteria from marine and non-marine sources are marked in blue and brown, respectively. The scale represents geological time in Gya.

Table S1 The calibration settings used in this study are presented in a format compatible with MCMCTree. The calibrations for the first-step dating analysis were adapted from our recent study (Zhang et al., 2023). In this step, only direct eukaryotic fossil-based calibrations were used. In the second-step dating analysis, additional eukaryotic fossils, as well as cyanobacterial and biomarker fossils, were incorporated. Notably, secondary calibrations derived from the first-step dating analysis were also used and approximated to either skew-t or skew-normal distributions (denoted as 'ST' or 'SN'). Ancestral nodes constrained by eukaryotic fossils are marked in red.

| Strategy | Node_ID | Calibrations in the first-step dating analysis | Calibrations in the second-step dating analysis | Note |
|---|---|---|---|---|
| Focal | Node_1 | NA | B(1.64,4.5,1e-300,1e-300) | Total Bacteroidetes/Chlorobia group (carotenoid biomarker) |
| | Node_2 | NA | B(1.6,4.5,1e-300,1e-300) | Total Nostocales group |
| | Node_3 | NA | B(1.7,4.5,1e-300,1e-300) | Total Pleurocapsales group |
| | Node_4 | B(0.947,1.891,1e-300,0.025) | SN(1.0,0.03,1.0) | Crown Chlorophyta group |
| | Node_5 | B(0.125,0.25,1e-300,0.025) | ST(0.25,0.01,3.39,11.41) | Crown Angiosperm group (Total Eudicots group) |
| | Node_6 | NA | B(0.308,0.509,1e-300,0.025) | Crown Spermatophyta group (Total Angiosperms group) |
| | Node_7 | B(0.45,0.509,1e-300,0.025) | SN(0.49,0.017,-0.95) | Crown Embryophyta group |
| | Node_8 | NA | SN(1.10,0.035,0.77) | NA |
| | Node_9 | B(0.947,1.891,1e-300,0.025) | SN(1.16,0.036,0.69) | Crown Viridiplantae group |
| | Node_10 | NA | B(0.08,1.891,1e-300,0.025) | Crown diatom group (Total pennate diatom group) |
| | Node_11 | NA | B(0.134,1.891,1e-300,0.025) | Total diatom group |
| | Node_12 | B(0.55,1.891,1e-300,0.025) | ST(0.91,0.038,0.031,33.93) | Total Florideophyceae group |
| | Node_13 | NA | ST(1.05,0.028,0.91,9.52) | NA |
| | Node_14 | B(1.2,1.891,1e-300,0.025) | ST(1.20,0.020,4753.67,3.57) | Crown Rhodophyta group |
| | Node_15 | NA | SN(1.54,0.042,0.47) | NA |
| | Node_16 | B(1.2,1.891,1e-300,0.025) | ST(1.51,0.065,2.10,68.65) | Crown Eukaryotes group (Archaeplastida) |
| | Node_17 | NA | B(2.32,4.5,1e-300,1e-300) | Total oxygenic Cyanobacteria |
| | Node_18 | NA | B(2.32,4.5,1e-300,1e-300) | Root |
| Topo-C1 | Node_1 | NA | B(1.64,4.5,1e-300,1e-300) | Total Bacteroidetes/Chlorobia group |
| | Node_2 | NA | B(1.6,4.5,1e-300,1e-300) | Total Nostocales group |
| | Node_3 | NA | B(1.7,4.5,1e-300,1e-300) | Total Pleurocapsales group |
| | Node_4 | B(0.947,1.891,1e-300,0.025) | ST(0.94,0.021,1697.91,3.11) | Crown Chlorophyta group |
| | Node_5 | B(0.125,0.25,1e-300,0.025) | ST(0.24,0.0063,0.50,4.02) | Crown Angiosperm group (Total Eudicots group) |
| | Node_6 | NA | B(0.308,0.509,1e-300,0.025) | Crown Spermatophyta group(Total Angiosperms group) |
| | Node_7 | B(0.45,0.509,1e-300,0.025) | SN(0.48,0.016,-0.015) | Crown Embryophyta group |
| | Node_8 | NA | ST(1.04,0.037,3.03,7.43) | NA |
| | Node_9 | B(0.947,1.891,1e-300,0.025) | ST(1.09,0.043,2.82,9.40) | Crown Viridiplantae group |
| | Node_10 | NA | ST(1.35,0.08,2.31,32.22) | NA |
| | Node_11 | NA | B(0.08, 1.9, 1.0e-300, 0.025) | Crown diatom group(Total pennate diatom group) |
| | Node_12 | NA | B(0.13, 1.9, 1.0e-300, 0.025) | Total diatom group |
| | Node_13 | B(0.55,1.891,1e-300,0.025) | ST(0.95, 0.086, 1.1, 17) | Total Florideophyceae group |
| | Node_14 | NA | ST(1.1, 0.098, 3.3, 9.6) | NA |
| | Node_15 | B(1.2,1.891,1e-300,0.025) | ST(1.2, 0.13, 5.0e+4, 2.0e+5) | Crown Rhodophyta group |
| | Node_16 | B(1.2,1.891,1e-300,0.025) | SN(1.6, 0.069, 0.58) | Crown Eukaryotes group (Archaeplastida) |
| | Node_17 | NA | B(2.3, 4.5, 1.0e-300, 1.0e-300) | Total oxygenic Cyanobacteria |
| | Node_18 | NA | B(2.3, 4.5, 1.0e-300, 1.0e-300) | Root |
| Topo-C2 | Node_1 | NA | B(1.6, 4.5, 1.0e-300, 1.0e-300) | Total Bacteroidetes/Chlorobia group |
| | Node_2 | NA | B(1.6, 4.5, 1.0e-300, 1.0e-300) | Total Nostocales group |
| | Node_3 | NA | B(1.7, 4.5, 1.0e-300, 1.0e-300) | Total Pleurocapsales group |
| | Node_4 | B(0.947,1.891,1e-300,0.025) | ST(0.95, 0.065, 1.0e+6, 1.3e+6) | Crown Chlorophyta group |
| | Node_5 | NA | ST(1.0, 0.068, 6.7, 23) | NA |
| | Node_6 | B(0.125,0.25,1e-300,0.025) | ST(0.25, 0.0059, -0.035, 3.9) | Crown Angiosperm group (Total Eudicots group) |
| | Node_7 | NA | B(0.31, 0.51, 1.0e-300, 0.025) | Crown Spermatophyta group(Total Angiosperms group) |
| | Node_8 | B(0.45,0.509,1e-300,0.025) | SN(0.48, 0.019, -0.95) | Crown Embryophyta group |
| | Node_9 | B(0.947,1.891,1e-300,0.025) | ST(1.1, 0.076, 4.4, 40) | Crown Viridiplantae group |
| | Node_10 | NA | B(0.08, 1.9, 1.0e-300, 0.025) | Crown diatom group(Total pennate diatom group) |
| | Node_11 | NA | B(0.13, 1.9, 1.0e-300, 0.025) | Total diatom group |
| | Node_12 | B(0.55,1.891,1e-300,0.025) | ST(0.92, 0.066, 0.63, 76) | Total Florideophyceae group |
| | Node_13 | NA | ST(1.0, 0.049, 1.3, 8.6) | NA |
| | Node_14 | B(1.2,1.891,1e-300,0.025) | ST(1.2, 0.048, 4.4e+4, 5.6) | Crown Rhodophyta group |
| | Node_15 | NA | SN(1.6, 0.064, 0.54) | NA |
| | Node_16 | B(1.2,1.891,1e-300,0.025) | SN(1.6, 0.070, 0.55) | Crown Eukaryotes group (Archaeplastida) |
| | Node_17 | NA | B(2.3, 4.5, 1.0e-300, 1.0e-300) | Total oxygenic Cyanobacteria |
| | Node_18 | NA | B(2.3, 4.5, 1.0e-300, 1.0e-300) | Root |

Table S2 The fitness test of the clock model for the gene families used in our dating analysis. The Bayes factor and posterior probability were calculated based on the marginal likelihood values for each candidate model using ‘mcmc3r’. IR and AR represent the independent clock model and the auto-correlated rate model, respectively.

| Gene_ID | Bayes_Factor | | Posterior model probability | |
|---|---|---|---|---|
| | IR | AR | IR | AR |
| NP_043188.1 | 1.78E-22 | 1.00E+00 | 1.78E-22 | 1.00E+00 |
| NP_043209.1 | 2.11E-16 | 1.00E+00 | 2.11E-16 | 1.00E+00 |
| NP_043214.1 | 1.00E+00 | 1.46E-56 | 1.00E+00 | 1.46E-56 |
| NP_043222.1 | 1.46E-13 | 1.00E+00 | 1.46E-13 | 1.00E+00 |
| NP_043226.1 | 0.1014564 | 1 | 0.0921111 | 0.9078889 |
| NP_043258.1 | 1.15E-24 | 1.00E+00 | 1.15E-24 | 1.00E+00 |
| NP_043267.1 | 2.96E-26 | 1.00E+00 | 2.96E-26 | 1.00E+00 |
| NP_043287.1 | 2.43E-26 | 1.00E+00 | 2.43E-26 | 1.00E+00 |

Table S3 The 84 secondary calibrations used to constrain the time prior of the flavobacterial 16S rRNA gene tree. The calibrations were obtained by fitting the posterior age estimates of flavobacteria, derived from the focal dating strategy (Fig. S4a), to a normal distribution. The paired "Taxon A" and "Taxon B" represent their last common ancestor for calibration. The Node_ID corresponds to those marked in Fig. S6. The two parameters of the normal distribution, i.e. the mean and standard deviation, are provided.

| Node_ID | Taxon_A | Taxon_B | Normal_mean | Normal_stddev |
|---|---|---|---|---|
| Node_1 | ref_Maribacter_thermophilus_HT7-2 | ref_Maribacter_cobaltidurans_B1 | 0.47948789 | 0.094654679 |
| Node_2 | ref_Maribacter_thermophilus_HT7-2 | ref_Maribacter_dokdonensis_MAR_2009_60 | 0.531252891 | 0.098748199 |
| Node_3 | ref_Maribacter_thermophilus_HT7-2 | ref_Maribacter_antarcticus_DSM_21422 | 0.554260529 | 0.101319599 |
| Node_4 | ref_Maribacter_thermophilus_HT7-2 | ref_Maribacter_vaceletii_DSM_25230 | 1.017602299 | 0.086695419 |
| Node_5 | ref_Maribacter_polysiphoniae_DSM_23514 | ref_Ulvibacterium_marinum_CCMM003 | 0.963288391 | 0.093442271 |
| Node_6 | ref_Maribacter_polysiphoniae_DSM_23514 | ref_Aggregatimonas_sangjinii_F202Z8 | 0.999906731 | 0.088325726 |
| Node_7 | ref_Maribacter_thermophilus_HT7-2 | ref_Aggregatimonas_sangjinii_F202Z8 | 1.044990158 | 0.084785565 |
| Node_8 | ref_Maribacter_thermophilus_HT7-2 | ref_Costertonia_aggregata_KCCM_42265 | 1.112367256 | 0.05823252 |
| Node_9 | ref_Euzebyella_marina_RN62 | ref_Pricia_antarctica_DSM_23421 | 0.917260803 | 0.090194329 |
| Node_10 | ref_Euzebyella_marina_RN62 | ref_Zobellia_galactanivorans | 0.996240141 | 0.07919388 |
| Node_11 | ref_Maribacter_thermophilus_HT7-2 | ref_Zobellia_galactanivorans | 1.122115048 | 0.057911646 |
| Node_12 | ref_Muricauda_ruestringensis_DSM_13258 | ref_Muricauda_aurantiaca_HME9304 | 0.544588509 | 0.087818696 |
| Node_13 | ref_Aureicoccus_marinus_SG-18 | ref_Muricauda_sp__DJ-13 | 0.64214241 | 0.103191913 |
| Node_14 | ref_Muricauda_ruestringensis_DSM_13258 | ref_Muricauda_sp__DJ-13 | 0.729919071 | 0.102261566 |
| Node_15 | ref_Robiginitalea_biformata_HTCC2501 | ref_Eudoraea_adriatica_DSM_19308 | 0.947808281 | 0.094675623 |
| Node_16 | ref_Muricauda_ruestringensis_DSM_13258 | ref_Eudoraea_adriatica_DSM_19308 | 1.127385121 | 0.06011625 |
| Node_17 | ref_Maribacter_thermophilus_HT7-2 | ref_Eudoraea_adriatica_DSM_19308 | 1.167142531 | 0.056938455 |
| Node_18 | ref_Arenibacter_latericius_DSM_15913 | ref_Arenibacter_algicola_SMS7 | 0.267423264 | 0.069900289 |
| Node_19 | ref_Arenibacter_latericius_DSM_15913 | ref_Cellulophaga_algicola_DSM_14237 | 0.641660791 | 0.099527799 |
| Node_20 | ref_Arenibacter_latericius_DSM_15913 | ref_Cellulophaga_lytica_DSM_7489 | 0.716014787 | 0.10054015 |
| Node_21 | ref_Maribacter_thermophilus_HT7-2 | ref_Cellulophaga_lytica_DSM_7489 | 1.203523777 | 0.056252094 |
| Node_22 | ref_Robertkochia_marina_CC-AMO-30D | ref_Robertkochia_solimangrovi_CL23 | 0.636051856 | 0.111801482 |
| Node_23 | ref_Robertkochia_marina_CC-AMO-30D | ref_Joostella_marina_DSM_19592 | 0.899789283 | 0.105567723 |
| Node_24 | ref_Robertkochia_marina_CC-AMO-30D | ref_Zhouia_spongiae_HN-Y44 | 1.009638123 | 0.093477327 |
| Node_25 | ref_Robertkochia_marina_CC-AMO-30D | ref_Abyssalbus_ytuae_MT3330 | 1.179924231 | 0.07242142 |
| Node_26 | ref_Capnocytophaga_ochracea_DSM_7271 | ref_Capnocytophaga_haemolytica_NCTC12947 | 0.533290864 | 0.108196363 |
| Node_27 | ref_Capnocytophaga_ochracea_DSM_7271 | ref_Imtechella_halotolerans_K1 | 1.08384298 | 0.092274046 |
| Node_28 | ref_Capnocytophaga_ochracea_DSM_7271 | ref_Sinomicrobium_kalidii_HD2P242 | 1.141728504 | 0.09294833 |
| Node_29 | ref_Robertkochia_marina_CC-AMO-30D | ref_Sinomicrobium_kalidii_HD2P242 | 1.275977446 | 0.068503408 |
| Node_30 | ref_Maribacter_thermophilus_HT7-2 | ref_Sinomicrobium_kalidii_HD2P242 | 1.399901748 | 0.060837175 |
| Node_31 | ref_Maribacter_thermophilus_HT7-2 | ref_Kordia_antarctica_IMCC3317 | 1.475531867 | 0.063841498 |
| Node_32 | ref_Psychroflexus_torquis_ATCC_700755 | ref_Psychroflexus_halocasei_DSM_23581 | 0.353748359 | 0.070698639 |
| Node_33 | ref_Psychroflexus_torquis_ATCC_700755 | ref_Mesohalobacter_halotolerans_WDS2C27 | 0.500821341 | 0.079458762 |
| Node_34 | ref_Psychroflexus_torquis_ATCC_700755 | ref_Croceibacter_atlanticus_HTCC2559 | 0.694446687 | 0.087220647 |
| Node_35 | ref_Zunongwangia_profunda_SM-A87 | ref_Salegentibacter_salegens_ACAM_48 | 0.593576729 | 0.08239251 |
| Node_36 | ref_Antarcticibacterium_flavum_JB01H24 | ref_Antarcticibacterium_arcticum_PAMC_28998 | 0.336303332 | 0.07872173 |
| Node_37 | ref_Zunongwangia_profunda_SM-A87 | ref_Antarcticibacterium_arcticum_PAMC_28998 | 0.655681359 | 0.084836053 |
| Node_38 | ref_Psychroflexus_torquis_ATCC_700755 | ref_Antarcticibacterium_arcticum_PAMC_28998 | 0.743970253 | 0.087292495 |
| Node_39 | ref_Aquimarina_agarilytica_ZC1 | ref_Aquimarina_brevivitae_DSM_17196 | 0.692505306 | 0.113427569 |
| Node_40 | ref_Psychroflexus_torquis_ATCC_700755 | ref_Aquimarina_brevivitae_DSM_17196 | 1.069005464 | 0.064621667 |
| Node_41 | ref_Nonlabens_marinus_S1-08 | ref_Nonlabens_spongiae_JCM_13191 | 0.399909503 | 0.113499012 |
| Node_42 | ref_Psychroflexus_torquis_ATCC_700755 | ref_Nonlabens_spongiae_JCM_13191 | 1.126098366 | 0.061035069 |
| Node_43 | ref_Dokdonia_donghaensis_DSW-1 | ref_Leeuwenhoekiella_palythoae_F5 | 0.731475513 | 0.133419427 |
| Node_44 | ref_Psychroflexus_torquis_ATCC_700755 | ref_Leeuwenhoekiella_palythoae_F5 | 1.156396769 | 0.059332919 |
| Node_45 | ref_Aequorivita_sublithincola_DSM_14238 | ref_Ulvibacter_antarcticus_DSM_23424 | 0.530927992 | 0.117577163 |
| Node_46 | ref_Aureitalea_marina_NBRC_107741 | ref_Rasiella_rasia_RR4-40 | 0.843238024 | 0.104815938 |
| Node_47 | ref_Aequorivita_sublithincola_DSM_14238 | ref_Rasiella_rasia_RR4-40 | 0.879607706 | 0.101178025 |
| Node_48 | ref_Psychroflexus_torquis_ATCC_700755 | ref_Rasiella_rasia_RR4-40 | 1.206159557 | 0.058513536 |
| Node_49 | ref_Algibacter_alginicilyticus_HZ22 | ref_Confluentibacter_flavum_3B_23 | 0.908684208 | 0.073466785 |
| Node_50 | ref_Algibacter_alginicilyticus_HZ22 | ref_Tamlana_carrageenivorans_UJ94 | 1.027186798 | 0.064814558 |
| Node_51 | ref_Formosa_agariphila_KMM_3901 | ref_Aurantibacter_aestuarii_KCTC_32269 | 1.101463667 | 0.062258032 |
| Node_52 | ref_Algibacter_alginicilyticus_HZ22 | ref_Aurantibacter_aestuarii_KCTC_32269 | 1.146336543 | 0.056803018 |
| Node_53 | ref_Gelidibacter_mesophilus_DSM_14095 | ref_Psychroserpens_burtonensis_DSM_12212 | 1.098048938 | 0.064096942 |
| Node_54 | ref_Gelidibacter_mesophilus_DSM_14095 | ref_Winogradskyella_sediminis_RHA_55 | 1.131732261 | 0.059086272 |
| Node_55 | ref_Algibacter_alginicilyticus_HZ22 | ref_Winogradskyella_sediminis_RHA_55 | 1.188212451 | 0.056060846 |
| Node_56 | ref_Psychroflexus_torquis_ATCC_700755 | ref_Winogradskyella_sediminis_RHA_55 | 1.484986036 | 0.063403283 |
| Node_57 | ref_Maribacter_thermophilus_HT7-2 | ref_Winogradskyella_sediminis_RHA_55 | 1.523687443 | 0.06515745 |
| Node_58 | ref_Flavobacterium_johnsoniae_UW101 | ref_Flavobacterium_branchiophilum_FL-15 | 0.355812512 | 0.085638792 |
| Node_59 | ref_Flavobacterium_soli_DSM_19725 | ref_Flavobacterium_noncentrifugens_CGMCC_1_10076 | 0.409507124 | 0.092998113 |
| Node_60 | ref_Flavobacterium_johnsoniae_UW101 | ref_Flavobacterium_noncentrifugens_CGMCC_1_10076 | 0.597843187 | 0.099650185 |
| Node_61 | ref_Flavobacterium_johnsoniae_UW101 | ref_Flavobacterium_aurantiibacter_TH167_C2897 | 0.773676542 | 0.10569293 |
| Node_62 | ref_Flavobacterium_johnsoniae_UW101 | ref_Flavobacterium_sangjuense_GS03 | 0.8452809 | 0.101536221 |
| Node_63 | ref_Flavobacterium_johnsoniae_UW101 | ref_Flavobacterium_azooxidireducens_IGB_4-14 | 0.947430338 | 0.095858938 |
| Node_64 | ref_Flavobacterium_covae_94-081 | ref_Flavobacterium_stagni_WWJ-16 | 1.012641007 | 0.092228436 |
| Node_65 | ref_Flavobacterium_johnsoniae_UW101 | ref_Flavobacterium_stagni_WWJ-16 | 1.111177312 | 0.080531287 |
| Node_66 | ref_Flavobacterium_johnsoniae_UW101 | ref_Flavobacterium_album_HYN0059 | 1.157037702 | 0.079522548 |
| Node_67 | ref_Flavobacterium_indicum_GPTSA100-9_= | ref_Flavobacterium_cyclinae_KSM-R2A25 | 0.918753573 | 0.138812588 |
| Node_68 | ref_Flavobacterium_johnsoniae_UW101 | ref_Flavobacterium_cyclinae_KSM-R2A25 | 1.233561903 | 0.082862519 |
| Node_69 | ref_Myroides_profundi_D25 | ref_Myroides_fluvii_CJ210 | 0.647476012 | 0.145811315 |
| Node_70 | ref_Flavobacterium_johnsoniae_UW101 | ref_Myroides_fluvii_CJ210 | 1.401244869 | 0.092308134 |
| Node_71 | ref_Maribacter_thermophilus_HT7-2 | ref_Myroides_fluvii_CJ210 | 1.6434846 | 0.074902893 |
| Node_72 | ref_Maribacter_thermophilus_HT7-2 | ref_Urechidicola_croceus_LPB0138 | 1.717082724 | 0.081703437 |
| Node_73 | ref_Maribacter_thermophilus_HT7-2 | ref_Ichthyobacterium_seriolicida_DNA | 1.910245532 | 0.097380617 |
| Node_74 | ref_Frigoriflavimonas_asaccharolytica_16F_Ga0416765_01 | ref_Chryseobacterium_manosquense_Marseille-Q2069 | 0.554092461 | 0.122409209 |
| Node_75 | ref_Frigoriflavimonas_asaccharolytica_16F_Ga0416765_01 | ref_Soonwooa_buanensis_DSM_22323 | 0.779873959 | 0.141497392 |
| Node_76 | ref_Epilithonimonas_vandammei_F5649 | ref_Chryseobacterium_aureum_17S1E7 | 0.728388521 | 0.140706088 |
| Node_77 | ref_Frigoriflavimonas_asaccharolytica_16F_Ga0416765_01 | ref_Chryseobacterium_aureum_17S1E7 | 0.826260513 | 0.143304933 |
| Node_78 | ref_Frigoriflavimonas_asaccharolytica_16F_Ga0416765_01 | ref_Cruoricaptor_ignavus_M1214 | 1.057969602 | 0.141855204 |
| Node_79 | ref_Frigoriflavimonas_asaccharolytica_16F_Ga0416765_01 | ref_Apibacter_raozihei_HY041 | 1.585043543 | 0.12518478 |
| Node_80 | ref_Empedobacter_falsenii_1681-1 | ref_Weeksella_virosa_NCTC11634 | 0.660238161 | 0.168141851 |
| Node_81 | ref_Empedobacter_falsenii_1681-1 | ref_Moheibacter_sediminis_CGMCC_1_12708 | 1.144214423 | 0.148709309 |
| Node_82 | ref_Empedobacter_falsenii_1681-1 | ref_Ornithobacterium_rhinotracheale_ORT-UMN_88 | 1.577242856 | 0.123484091 |
| Node_83 | ref_Frigoriflavimonas_asaccharolytica_16F_Ga0416765_01 | ref_Ornithobacterium_rhinotracheale_ORT-UMN_88 | 1.697139221 | 0.117207334 |
| Node_84 | ref_Maribacter_thermophilus_HT7-2 | ref_Ornithobacterium_rhinotracheale_ORT-UMN_88 | 1.983592817 | 0.10203302 |
